# Polycomb repressive complex 1 shapes the nucleosome landscape but not accessibility at target genes

**DOI:** 10.1101/280305

**Authors:** Hamish W King, Nadezda A Fursova, Neil P Blackledge, Robert J Klose

## Abstract

Polycomb group (PcG) proteins are transcriptional repressors that play important roles regulating gene expression during animal development. In *vitro* experiments have shown that PcG protein complexes can compact chromatin to limit the activity of chromatin remodelling enzymes and access of the transcriptional machinery to DNA. In fitting with these ideas, gene promoters associated with PcG proteins have been reported to be less accessible than other gene promoters. However, it remains largely untested *in vivo* whether PcG proteins define chromatin accessibility or other chromatin features. To address this important question, we examine the chromatin accessibility and nucleosome landscape at PcG protein-bound promoters in mouse embryonic stem cells using the assay for transposase accessible chromatin (ATAC)-seq. Combined with genetic ablation strategies, we unexpectedly discover that although PcG protein-occupied gene promoters exhibit reduced accessibility, this does not rely on PcG proteins. Instead, the Polycomb repressive complex 1 (PRC1) appears to play a unique role in driving elevated nucleosome occupancy and decreased nucleosomal spacing in Polycomb chromatin domains. Our new genome-scale observations argue, in contrast to the prevailing view, that PcG proteins do not significantly affect chromatin accessibility and highlight an underappreciated complexity in the relationship between chromatin accessibility, the nucleosome landscape and PcG-mediated transcriptional repression.

## INTRODUCTION

In eukaryotic cells, DNA is wrapped around histone octamers to form nucleosomes and chromatin (Kornberg and Lorch 1999). Chromatin functions to organise the DNA of large eukaryotic genomes into the relatively small confines of the nucleus. The position of nucleosomes on DNA and the organisation of nucleosomes into higher order chromatin structures also plays major roles in gene regulation (Kouzarides 2007; Li et al. 2007). For example, nucleosomes can occlude sequence-specific transcription factors and the transcriptional machinery from accessing the DNA sequence, thus regulating their activity (Kornberg and Lorch 1999; Li et al. 2007; Jiang and Pugh 2009). This can be overcome through the eviction, repositioning or destabilisation of nucleosomes to create chromatin states that are more accessible to *trans*-acting factors (Henikoff 2008; Jiang and Pugh 2009). Accessible chromatin is therefore a characteristic feature of gene regulatory elements including gene promoters and enhancers (Boyle et al. 2008; Song et al. 2011; Thurman et al. 2012). The formation and maintenance of accessible chromatin states appears to be highly regulated and accessibility is often related to post-translational modification of histones associated with gene regulatory elements. By extension, it has been proposed that chromatin-modifying systems and their associated activities may help to define accessibility at these important regulatory sites.

In animals, Polycomb group (PcG) proteins play central roles in developmental gene regulation. This diverse group of proteins form large multi-protein complexes that bind gene regulatory elements and modify chromatin to establish what is thought to be a transcriptionally repressive chromatin state (Muller and Verrijzer 2009; Di Croce and Helin 2013). PcG proteins generally exist in one of two multi-protein complexes, known as Polycomb repressive complexes 1 and 2 (PRC1 and PRC2). PRC1 complexes, through their catalytic subunit, RING1 (also known as RING1A) or RNF2 (also known as RING1B), mono-ubiquitylate histone H2A at lysine 119 (H2AK119ub1), while PRC2 methylates histone H3 on lysine 27 (H3K27me3). In vertebrates, PRC1 and PRC2 are targeted by various mechanisms to gene promoters, particularly those associated with non-methylated CpG islands (CGIs) (Bracken and Helin 2009; Simon and Kingston 2013; Blackledge et al. 2015). The occupancy and activity of PcG complexes at CGI gene promoters is typically associated with low or undetectable transcriptional activity. Removal of PcG complexes can lead to the abnormal transcription of PcG-occupied genes (Boyer et al. 2006; Bracken et al. 2006; Endoh et al. 2008; Leeb et al. 2010; Blackledge et al. 2014). Many of these inappropriately activated genes are associated with embryonic development and their precocious expression during embryogenesis could possibly explain the embryonic lethal phenotypes observed in PcG mutant mice. However, the mechanisms by which PcG complexes achieve transcriptional repression, and how this relates to their activities on chromatin, remain poorly understood.

PcG complexes are thought to repress transcription through the biochemical compaction of chromatin and the creation of inaccessible chromatin at PcG-occupied promoters. This is based in part on *in vitro* characterisation of reconstituted *Drosophila* and mammalian PRC1 complexes which were capable of compacting nucleosomal arrays and inhibiting the activity of nucleosome remodelling complexes (Francis et al. 2004; Trojer et al. 2007; Grau et al. 2011; Trojer et al. 2011). These biochemical studies supported a model whereby PcG complexes, particularly PRC1, compact chromatin to create a transcriptionally repressive chromatin state at PcG target sites. In fitting with these *in vitro* activities, *in vivo* PcG-occupied promoters exhibit reduced sensitivity to nuclease digestion when compared to gene promoters lacking PcG complexes (Bell et al. 2010; Calabrese et al. 2012; Kelly et al. 2012; Beck et al. 2014; Deaton et al. 2016). Furthermore, PcG target sites are also more refractory to transcription factor and polymerase binding (Zink and Paro 1995; McCall and Bender 1996; Fitzgerald and Bender 2001). Together, these studies have suggested that DNA within PcG complex-occupied chromatin is less accessible to *trans*-acting factors, consistent with biochemical activities that act locally to compact nucleosomes. However, despite the correlation between PcG protein occupancy and reduced accessibility, and the widespread view that PcG complexes create inaccessible chromatin, genome-scale analyses of whether PcG complexes directly influence chromatin accessibility *in vivo* remain limited. We therefore have a poor understanding of how chromatin organisation is achieved at PcG target sites and how this relates to their repressed transcriptional state.

To address these fundamental questions, we have interrogated chromatin at PcG-occupied gene promoters at the genome-scale using the assay for transposase accessible chromatin coupled with massively parallel sequencing (ATAC-seq) in mouse embryonic stem cells. We observe that PcG-occupied promoters exhibit reduced chromatin accessibility, elevated nucleosome occupancy and shorter inter-dyad distances compared to other gene promoters. Deletion of the PRC1 and PRC2 complexes individually, or in combination, uncovered a role for PRC1 in subtly shaping several features of the nucleosome landscape at PcG-occupied promoters, including nucleosome occupancy and spacing, but unexpectedly not accessibility. The regulation of the nucleosome landscape by PRC1 appeared to be linked to its transcriptionally repressive function, as genes activated after loss of PRC1 showed reversion towards a nucleosome landscape consistent with non-PcG occupied gene promoters. Together, these observations demonstrate a role for PRC1 in regulating the nucleosome landscape, but not accessibility, at PcG-occupied gene promoters and suggest that some of the prevailing models used to describe how PcG complexes control gene expression require re-evaluation.

## RESULTS

### Polycomb-occupied promoters show reduced accessibility

Previous studies have demonstrated that PcG proteins and their associated complexes are capable of compacting nucleosomal arrays *in vitro* (Francis et al. 2004; Trojer et al. 2007; Grau et al. 2011; Trojer et al. 2011), while *in vivo* studies have revealed reduced chromatin accessibility at PcG-occupied promoters compared to PcG-free promoters (Bell et al. 2010; Calabrese et al. 2012; Kelly et al. 2012; Beck et al. 2014; Deaton et al. 2016). These nucleosome-based features are distinct from other descriptions of PcG-dependent compaction which occur on the order of ~100-1000 kilobases and are more likely to reflect long-range chromatin interactions and higher order chromatin structures (Eskeland et al. 2010; Schoenfelder et al. 2015; Cruz-Molina et al. 2017; Kundu et al. 2017). To understand whether PcG complexes might indeed regulate the chromatin- and nucleosome-based landscape at gene promoters in living cells, we set out to carefully compare promoters occupied by PcG complexes and those which lack PcG proteins in mouse embryonic stem cells (ESCs) where PcG systems have been extensively studied. To achieve this, we first identified PcG-occupied promoters based on chromatin immunoprecipitation followed by massively parallel sequencing (ChIP-seq) for the PRC1 subunit RNF2 and the PRC2 subunit SUZ12 in mouse ESCs (Figure 1A). In fitting with previous studies that suggest CpG islands (CGIs) represent an important PcG recruitment site (Mendenhall et al. 2010; Farcas et al. 2012; He et al. 2013; Wu et al. 2013), we observed that 98.4% of PcG-occupied transcription start sites (TSS) are also marked by the presence of an experimentally-identified, non-methylated CGI (Figure 1B) (Long et al. 2013). Given the substantial differences between CGI and non-CGI chromatin (Blackledge and Klose 2011; Deaton and Bird 2011), we chose to focus our subsequent analysis to CGI-associated gene promoters. Having identified a high-confidence set of PcG-occupied promoters, we then examined whether PcG-occupied gene promoters were associated with chromatin that differed in any way from non-PcG promoters.

**Figure 1.**
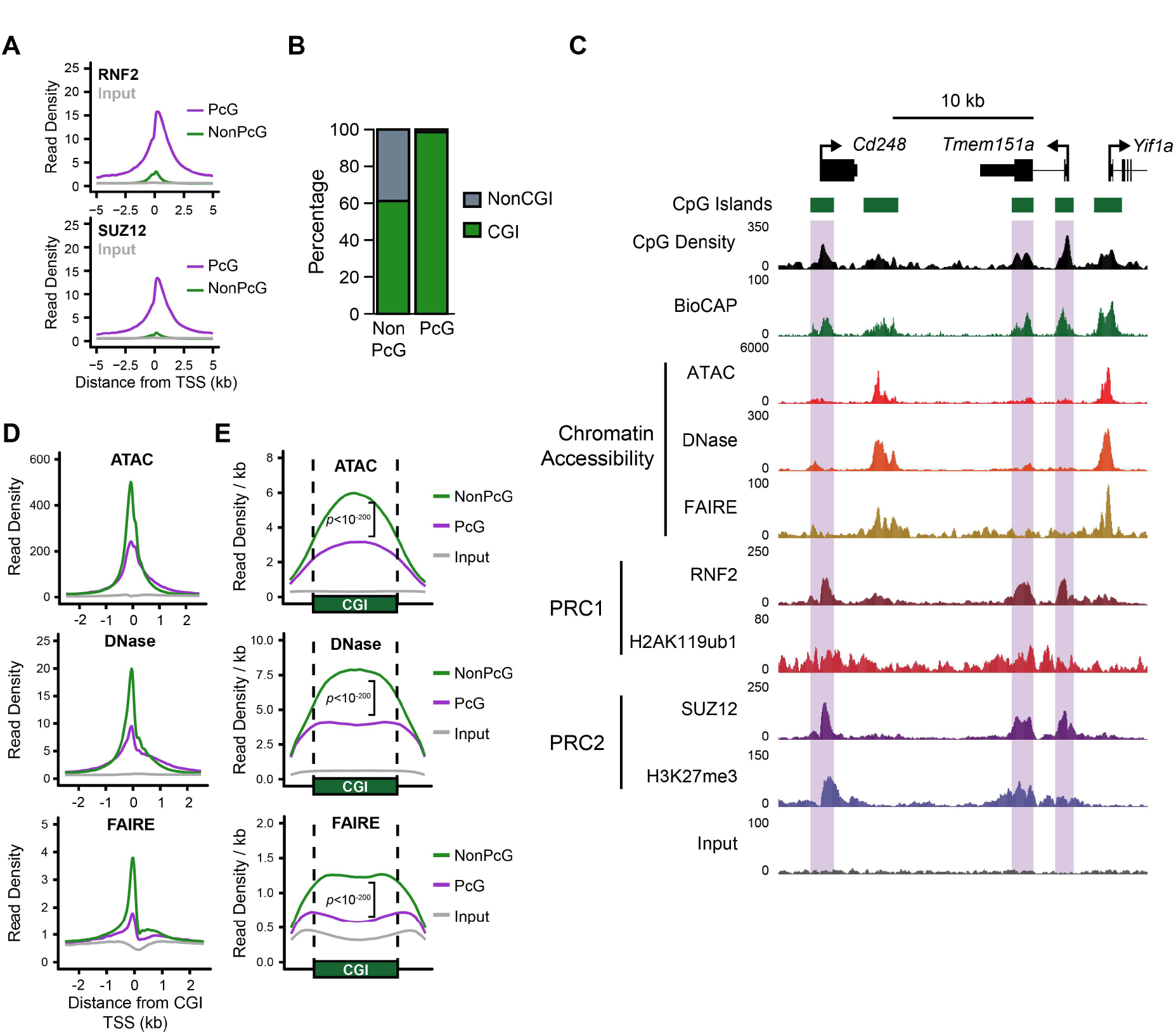
Polycomb-occupied promoters exhibit reduced chromatin accessibility compared to Polycomb-free promoters. A) A metaplot analysis comparing RNF2 (PRC1; upper panel) and SUZ12 (PRC2; lower panel) ChIP-seq signal at Polycomb (PcG)-occupied promoters or PcG-free (non-PcG) promoters in mouse embryonic stem cells (ESCs), centred on transcription start sites (TSS). B) A comparison of the percentage of PcG and non-PcG TSS (±500bp) that overlap with experimentally-identified non-methylated CpG islands (CGIs). C) A genome screenshot of several PcG-occupied promoters (highlighted in purple boxes) profiling three measures of chromatin accessibility, ATAC-seq, DNase-seq and FAIRE-seq. CpG density and non-methylated DNA (BioCAP), in addition to PRC1 and PRC2 ChIP-seq, are included for reference. D) A metaplot analysis comparing ATAC-seq, DNase-seq and FAIRE-seq signal at PcG-occupied (n = 4020) or PcG-free (n = 10251) CGI promoters, centred on TSS. Input for ATAC-seq and DNase-seq represents digestion of naked genomic DNA by Tn5 or DNase I respectively. E) A metaplot analysis at CGI intervals (±20%) for CGI-positive TSS with (PcG) or without (NonPcG) for ATAC-seq, DNase-seq and FAIRE-seq signal, normalised to CGI interval size. *p* values represent comparison of reads per kilobase per million (RPKM) at PcG-bound CGI promoter intervals compared to non-PcG CGI promoters.

We considered three different measures of chromatin accessibility in mouse embryonic stem cells: the assay for transposase accessible chromatin (ATAC-seq), DNase I hypersensitivity (DNase-seq) and formaldehyde-assisted isolation of regulatory elements (FAIRE-seq). ATAC-seq and DNase-seq measure accessibility by interrogating the digestion frequency of chromatin by Tn5 transposase or DNase I respectively (Cockerill 2011; Buenrostro et al. 2013). Alternatively, FAIRE-seq uses a biochemical approach to purify DNA fragments that are not physically bound by proteins (e.g. nucleosomes or transcription factors), providing a complimentary measure of whether a genomic locus exists in an accessible state (Giresi et al. 2007). Using these measurements, we compared chromatin accessibility at promoters with or without PcG complex occupancy (Figure 1C-E). Visual examination of several promoters occupied by PcG proteins clearly demonstrated that they had reduced accessibility when compared to neighbouring PcG-free promoters (Figure 1C). Indeed, a genome-wide analysis confirmed that PcG-occupied promoters exhibited significantly lower levels of accessibility than PcG-free promoters (Figure 1D), confirming and extending previous observations in both *Drosophila* and mammalian cells (Zink and Paro 1995; McCall and Bender 1996; Boivin and Dura 1998; Fitzgerald and Bender 2001; Bell et al. 2010; Calabrese et al. 2012; Beck et al. 2014; Deaton et al. 2016). This difference in accessibility was not limited to the TSS itself, but instead occurred across the entire breadth of the CGI and its associated PcG chromatin domain (Figure 1E). Furthermore, PcG complex occupancy was associated with reduced chromatin accessibility when expression-matched PcG and non-PcG promoters were compared (Supplemental Fig. S1A). We also examined the relationship between PcG chromatin domains and promoter accessibility in other mouse tissues and cell lines. We found that reduced chromatin accessibility at PcG promoters was widespread (Supplemental Fig. S1B-C). Although PcG complex enrichment is usually associated with gene promoters in mammalian cells, we also examined the subset of distal elements bound by PcG complexes and compared them to distal elements that lacked PcG complexes (Supplemental Fig. S2A-B). In contrast to gene promoters, PcG-bound distal elements showed little difference in their accessibility when compared to distal elements without PcG binding. This suggested that PcG complexes may specifically limit chromatin accessibility at gene promoters.

### Elevated occupancy and closer spacing of nucleosomes at PcG-occupied promoters

PcG promoters have been proposed to exist in a more nucleosome-enriched state compared to non-PcG promoters in mammalian cells (Kelly et al. 2012; West et al. 2014). We were therefore keen to explore in more detail the nucleosome landscape at PcG-occupied gene promoters in our ATAC-seq experiments. To achieve this, we extracted nucleosome occupancy and positioning data using the NucleoATAC approach (Schep et al. 2015) (Figure 2A) and compared PcG-occupied promoters and PcG-free promoters in mouse embryonic stem cells (Figure 2B-F). Here we use the term nucleosome occupancy to describe the observed level of mono-nucleosome signal at defined nucleosome dyad centres, while nucleosome spacing refers to the distance between identified nucleosome positions (Figure 2A; see Methods for more details). Our analysis revealed elevated nucleosome occupancy at PcG promoters in agreement with previous observations (Figure 2B-D) (Kelly et al. 2012; West et al. 2014). One of the proposed functions of PcG complexes is to compact nucleosomal arrays (Francis et al. 2004; Trojer et al. 2007; Grau et al. 2011; Trojer et al. 2011). We therefore examined the spacing between nucleosomes at PcG-occupied promoters and observed that nucleosomes at PcG-occupied promoters exhibited shorter inter-dyad distances and their positions were less well-defined when compared to nucleosomes found at PcG-free promoters (Figure 2C,E-F). Elevated nucleosome occupancy and less well-positioned nucleosomes was clearly evident at PcG-bound promoters across all expression quantiles (Supplemental Fig. S3A), as well as at PcG-bound distal regulatory elements (Supplemental Fig. S2B). Together, these observations indicate that PcG target sites in ESCs exist in a nucleosome-rich state with more closely spaced nucleosomes than PcG-free regions.

**Figure 2.**
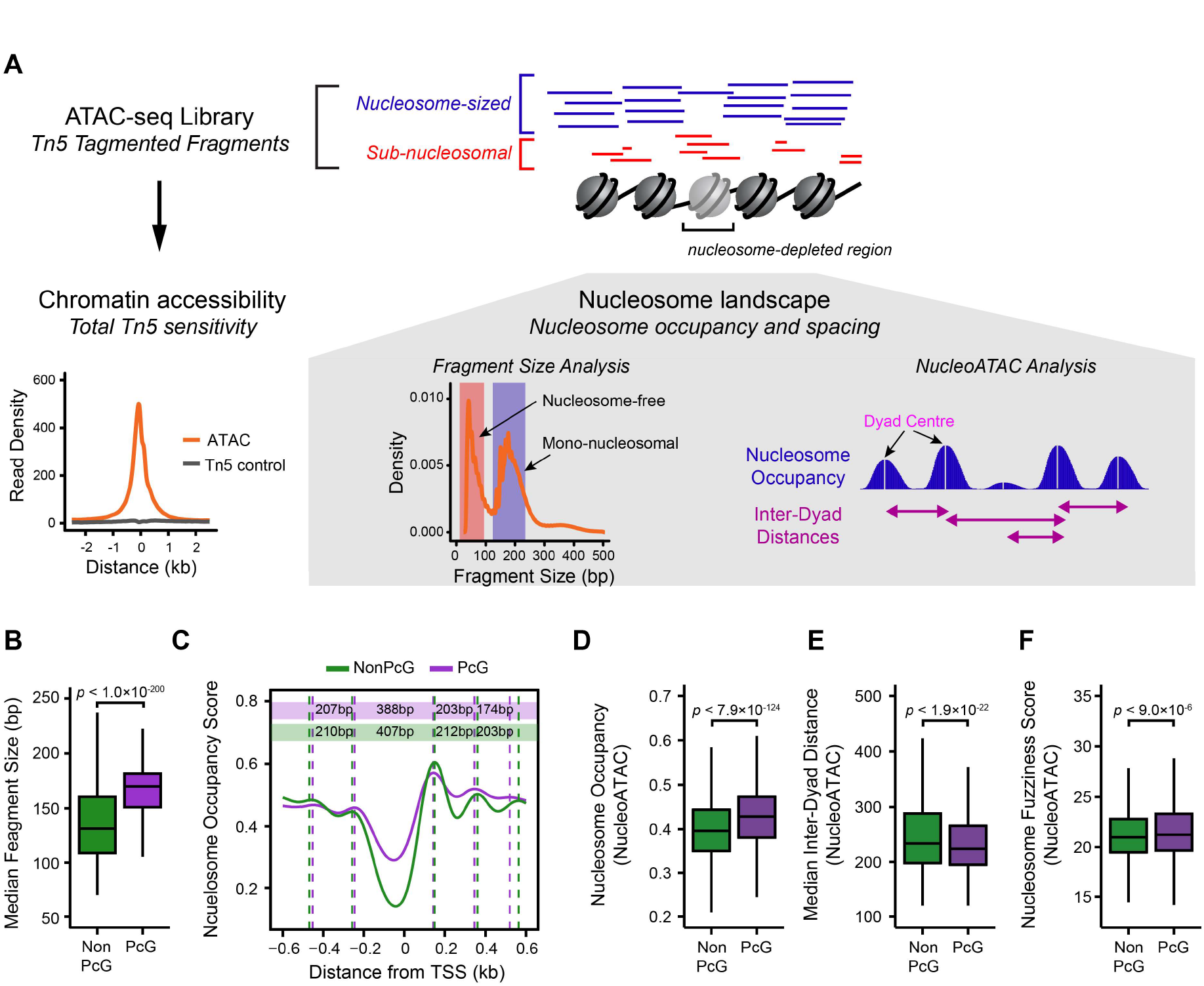
Characterisation of the nucleosome landscape at Polycomb-occupied promoters. A) A schematic detailing the approach to analyse nucleosome landscape features from ATAC-seq data. The cleavage of Tn5 hypersensitive DNA (accessible DNA) by Tn5 generates DNA fragments that broadly reflect either mono-nucleosomal fragments (blue) or nucleosome-free fragments (red). The total count of fragments represents total chromatin accessibility at a given loci, while the fragment size distribution allows the examination of qualitative features of Tn5 sensitivity, such as nucleosome occupancy or positioning using either the median fragment size for a gene promoter or the quantification of nucleosome occupancy signal using the software package NucleoATAC (Schep et al. 2015). After identifying nucleosome positions using NucleoATAC, individual nucleosome dyad centres can then be identified and the distance between neighbouring dyad centres can be calculated. B) A boxplot comparing the median ATAC-seq fragment sizes for PcG-occupied (n = 4020) or non-PcG (n = 10251) CGI promoters. PcG-occupied promoters tend to have larger fragment sizes consistent with an enrichment for nucleosomal-sized fragments. C) A metaplot for PcG-occupied or PcG-free promoters depicting nucleosome occupancy signal extracted from ATAC-seq data using NucleoATAC, centred on TSS. The average dyad centre for each nucleosome position is marked by dashed lines, and the distance between each nucleosome position is included in the coloured rectangles (Purple = PcG; Green = NonPcG). D) A boxplot comparing the NucleoATAC-derived nucleosome occupancy score within PcG-occupied or PcG-free promoters. E) A boxplot comparing the median inter-dyad distances within PcG-occupied or PcG-free promoters. Distances were calculated between the centres of neighbouring dyad positions identified by NucleoATAC. F) A boxplot comparing the median fuzziness score for nucleosomes identified by NucleoATAC within PcG-occupied or PcG-free promoters.

### Deletion of PRC1, but not PRC2, results in altered nucleosome occupancy and spacing without changes in chromatin accessibility

Our characterisation of the chromatin landscape at PcG-occupied promoters is consistent with previous reports implicating PcG complexes in the compaction of nucleosome arrays to create inaccessible chromatin. However, whether PcG complexes themselves define these features *in vivo* has yet to be interrogated satisfactorily. We therefore set out to examine the chromatin landscape of PcG-occupied promoters in cells lacking normal PcG complex activity (Figure 3). To achieve this, we exploited mouse ESC lines to ablate either PRC1 or PRC2. We used *Ring1*^-/-^;*Rnf2^fl/fl^* conditional ESCs (Endoh et al. 2008) in which the addition of tamoxifen leads to *Rnf2* deletion and the creation of PRC1-null cells (Figure 3A). As expected, treatment of this cell line with tamoxifen was sufficient to remove RNF2 protein and PRC1-deposited H2AK119ub1 (Figure 3B). Alternatively, the PRC2 core complex was removed using an EED conditional knockout cell line (*Eed^-/-^*;*Eed4. TG^DOX^*) which expresses a doxycycline-sensitive *Eed4* transgene (*Eed4^TG^*) in an *Eed^-/-^* background (Figure 3C; Ura et al. 2008). In the presence of doxycycline, *Eed4*^TG^ is not expressed, leading to loss of EED expression, destabilisation of the core PRC2 complex (Ura et al. 2008; Tavares et al. 2012) and loss of H3K27me3 (Figure 3D). We performed ATAC-seq in the *Ring1*^-/-^;*Rnf2^fl/fl^* and *Eed*^-/-^;*Eed4.TG^DOX^* ESCs in order to understand whether PRC1 or PRC2 are responsible for the chromatin features associated with PcG occupancy in mouse ESCs. Initially we considered two PcG-occupied genes, *Lhx9* and *Ovol1*, with low chromatin accessibility at their promoters in wild type ESCs and examined their accessibility in the PRC1- or PRC2-null state (Figure 3E). Surprisingly, there was no apparent change in chromatin accessibility in the absence of either PRC1 or PRC2 at these loci. We then extended this analysis across all PcG-occupied promoters. Again, we did not identify any significant changes in chromatin accessibility following deletion of either PRC1 or PRC2 (Figure 3F-G; Supplemental Fig. S4), in agreement with a previous study examining chromatin accessibility in the *Ring1*^-/-^;*Rnf2^fl/fl^* ESCs (Hodges et al. 2018). This was unexpected given the previously observed biochemical activities of PcG complexes, therefore revealing that deletion of PRC1 or PRC2 does not influence chromatin accessibility at PcG-occupied gene promoters in ESCs. To determine whether this was also the case in other cell types, we examined chromatin accessibility in a *Ring1*^-/-^;*Rnf2^fl/fl^* conditional mouse embryonic fibroblast cell line by ATAC-seq (Supplemental Fig. S5A-C). In wild type fibroblasts PcG-occupied promoters exhibited lower levels of chromatin accessibility compared to PcG-free promoters. In agreement with our analysis of PRC1-null ESCs, this disparity between chromatin accessibility of PcG-bound promoters and PcG-free promoters remained after deletion of PRC1 in the *Ring1*^-/-^;*Rnf2^fl/fl^* fibroblasts (Supplemental Fig S5C), although we did observe modest and non-specific increases in accessibility more generally. Therefore, we conclude that although PcG-bound promoters are associated with reduced accessibility compared to PcG-free promoters, PcG systems do not directly create this lack of accessibility.

**Figure 3.**
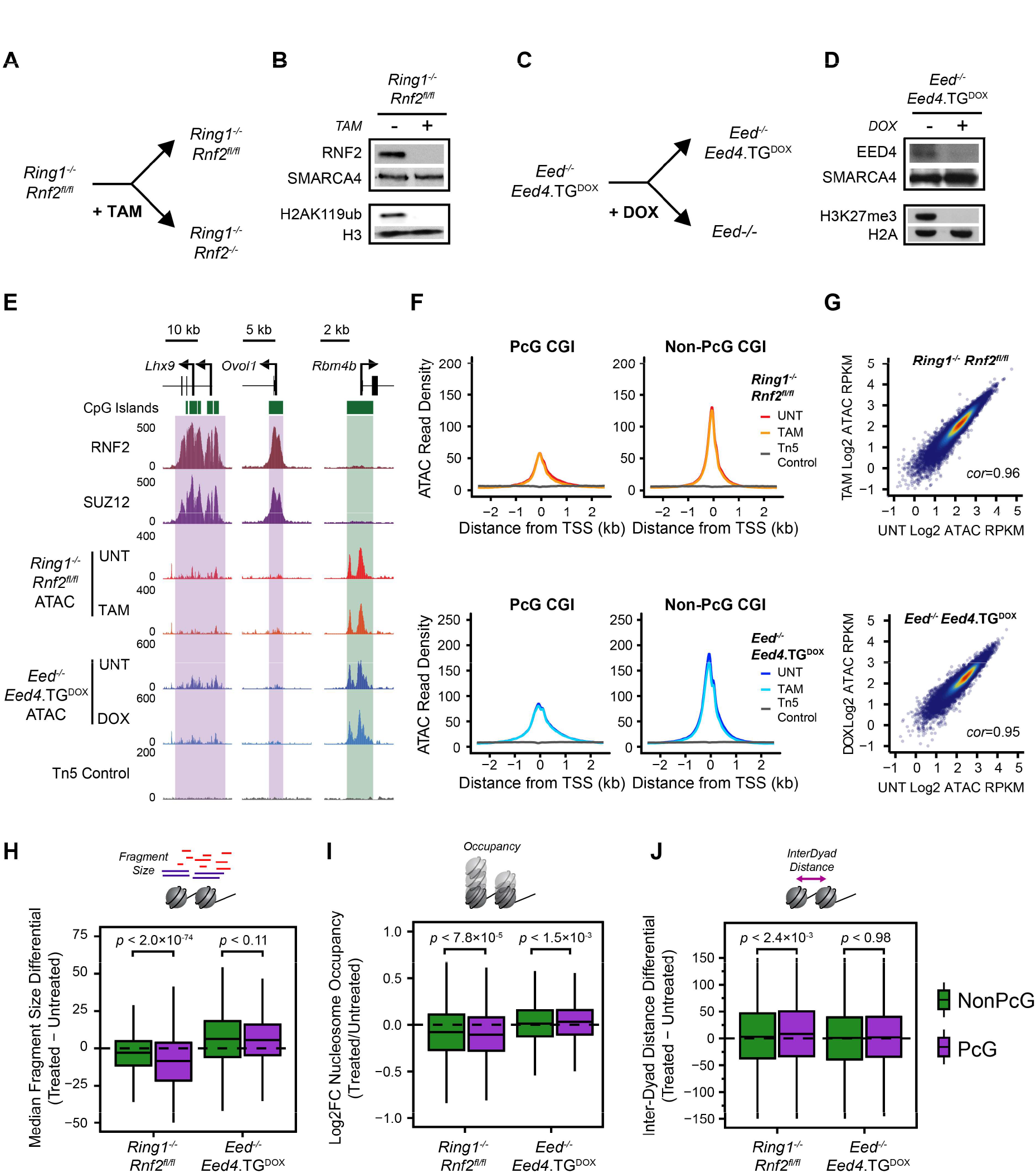
PRC1 contributes towards nucleosome spacing and occupancy but not chromatin accessibility. A) A schematic depicting the treatment of *Ring1*^-/-^;*Rnf2^fl/fl^* ESCs with 4-hydroxytamoxifen (TAM) to generate PRC1-null ESCs. B) A Western blot analysis of untreated and TAM-treated *Ring1*^-/-^;*Rnf2^fl/fl^* ESCs for RNF2 and H2AK119ub1. C) A schematic depicting the treatment of *Eed*^-/-^;*Eed4.TG^DOX^* ESCs with doxycycline (DOX) to generate PRC2-null ESCs. D) A Western blot analysis of untreated and DOX-treated *Eed^-/-^*;*Eed4.TG^DOX^* ESCs for EED and H3K27me3. E) Genome screenshots of chromatin accessibility, as measured by ATAC-seq, at two PcG-occupied promoters (purple boxes) and one PcG-free promoter (green box) before and after conditional deletion of PRC1 or PRC2 from mouse ESCS. F) A metaplot analysis for *Ring1*^-/-^;*Rnf2^fl/fl^* and *Eed*^-/-^;*Eed4.TG^DOX^* ATAC-seq before and after TAM or DOX treatment respectively, at PcG-occupied (*n* = 4020) or non-PcG CGI promoters (*n* = 10251), centred on TSS. G) A scatterplot analysis comparing untreated and treated reads per kilobase per million (RPKM) for *Ring1*^-/-^;*Rnf2^fl/fl^* and *Eed*^-/-^;*Eed4.TG^DOX^* ATAC-seq at all CGI promoters. H) A boxplot comparing the change in median Tn5-tagmented fragment sizes for PcG and non-PcG CGI promoters in *Ring1*^-/-^;*Rnf2^fl/fl^* and *Eed*^-/-^;*Eed4.TG^DOX^* ATAC-seq datasets before and after TAM or DOX treatment respectively. A decrease in median fragment size reflects a shift towards a more nucleosome-free state. I) A boxplot quantifying the log2 fold change (log2FC) in NucleoATAC-derived nucleosome occupancy for PcG and non-PcG CGI promoters in *Ring1*^-/-^;*Rnf2^fl/fl^* and *Eed*^-/-^;*Eed4.TG^DOX^* ATAC-seq datasets before and after TAM or DOX treatment respectively. J) A boxplot comparing the difference in median inter-dyad distances for PcG and non-PcG CGI promoters in *Ring1*^-/-^;*Rnf2^fl/fl^* and *Eed*^-/-^;*Eed4.TG^DOX^* ATAC-seq datasets before and after TAM or DOX treatment respectively.

Given that PcG-occupied gene promoters also show elevated nucleosome occupancy and closer nucleosome spacing (Figure 2), we were keen to examine whether these nucleosome features might be altered in the absence of either PRC1 or PRC2 (Figure 3H-J). These analyses revealed that the deletion of PRC1, but not PRC2, resulted in reductions in the mono-nucleosome sized fragments and NucleoATAC-derived nucleosome occupancy scores (Figure 3H-I, Supplemental Fig. S6A) coupled with subtle increases in inter-nucleosomal spacing (Figure 3J) at PcG-occupied promoters. We also performed MNase-seq as an alternative measurement of nucleosome positioning and occupancy in the *Ring1*^-/-^;*Rnf2^fl/fl^* ESCs (Supplemental Fig. S6B-D) and examined ATAC-derived nucleosome features in *Ring1*^-/-^;*Rnf2^fl/fl^* mouse embryonic fibroblasts (Supplemental Fig. S5D). These analyses revealed similar reductions in nucleosome occupancy and altered nucleosome spacing in the absence of PRC1, although this was less apparent in mouse embryonic fibroblasts. Importantly, these effects were not observed or considerably less dramatic at non-PcG promoters, indicating that this effect was specific to the promoters occupied by PcG complexes. To our knowledge, these observations demonstrate for the first time *in vivo* that PRC1 can influence the nucleosome landscape by altering nucleosome occupancy and spacing, albeit modestly, in a way that does not appear to define overall accessibility at the gene promoters. This suggests that these features are not directly coupled at PcG-occupied gene promoters.

### PcG complexes do not function redundantly to shape the chromatin landscape at PcG-occupied promoters

Previous studies have identified some instances of redundancy between the activity and function of PRC1 and PRC2 (Leeb et al. 2010). Furthermore, deletion of PRC1 results in widespread reductions, but not complete loss, of PRC2 at PcG-occupied promoters (Blackledge et al. 2014), and *vice versa* in PRC2-null cells (Tavares et al. 2012). As such it seemed possible that redundancy between PRC1 and PRC2 could potentially mask any effects on chromatin accessibility or more profound effects on nucleosome features at PcG-occupied promoters. We therefore sought to develop a cell culture system in which we could remove both PRC1 and PRC2. Previous reports have established that mouse ESCs lacking PRC2 are viable and can be maintained in culture (Boyer et al. 2006; Chamberlain et al. 2008; Shen et al. 2008; Leeb et al. 2010), while cells lacking PRC1 differentiate and are unable to be maintained as pluripotent cells (Stock et al. 2007; Endoh et al. 2008). Therefore, we constitutively deleted *Eed* (EED) in the *Ring1*^-/-^;*Rnf2^fl/fl^* conditional ESCs using CRISPR/Cas9-mediated gene editing (Figure 4A) and confirmed loss of EED protein levels by Western blotting (Figure 4B). Treatment of *Ring1*^-/-^;*Rnf2^fl/fl^*;*Eed*^-/-^ ESCs with tamoxifen resulted in the complete loss of PRC1, effectively removing both PcG complexes (Figure 4B). We then examined the chromatin landscape at PcG-occupied gene promoters by performing ATAC-seq on the *Ring1*^-/-^;*Rnf2^fl/fl^*;*Eed*^-/-^ ESCs with or without tamoxifen treatment. Even in the absence of both PRC1 and PRC2, there were no increases in accessibility at either individual PcG-occupied promoters (Figure 4C) or genome-wide (Figure 4D), similar to our observation in lines with deletion of PRC1 or PRC2 individually. However, consistent with a role for PRC1, but not PRC2, in modulating nucleosome spacing and occupancy at PcG-occupied promoters, we observed a shift towards nucleosome-free DNA and reduced nucleosome occupancy (Figure 4E-G), as well as increased inter-dyad spacing (Figure 4H), at PcG-bound promoters only in the PRC1- and PRC1/2-null cells. The effects in PRC1/2-null cells were similar to those observed in cells lacking only PRC1 (Figure 4E-H), suggesting little if any contribution of PRC2 to the regulation of nucleosome occupancy and spacing at PcG target sites, although we cannot exclude the possibility that this may reflect an adaptation of the nucleosome landscape in these cells due to constitutive loss of PRC2. Importantly, the fact that neither PRC1 or PRC2 appear to be responsible for limiting chromatin accessibility of PcG-occupied gene promoters in ESCs suggests that other pathways or processes must determine the reduced accessibility at these sites (see Discussion).

**Figure 4.**
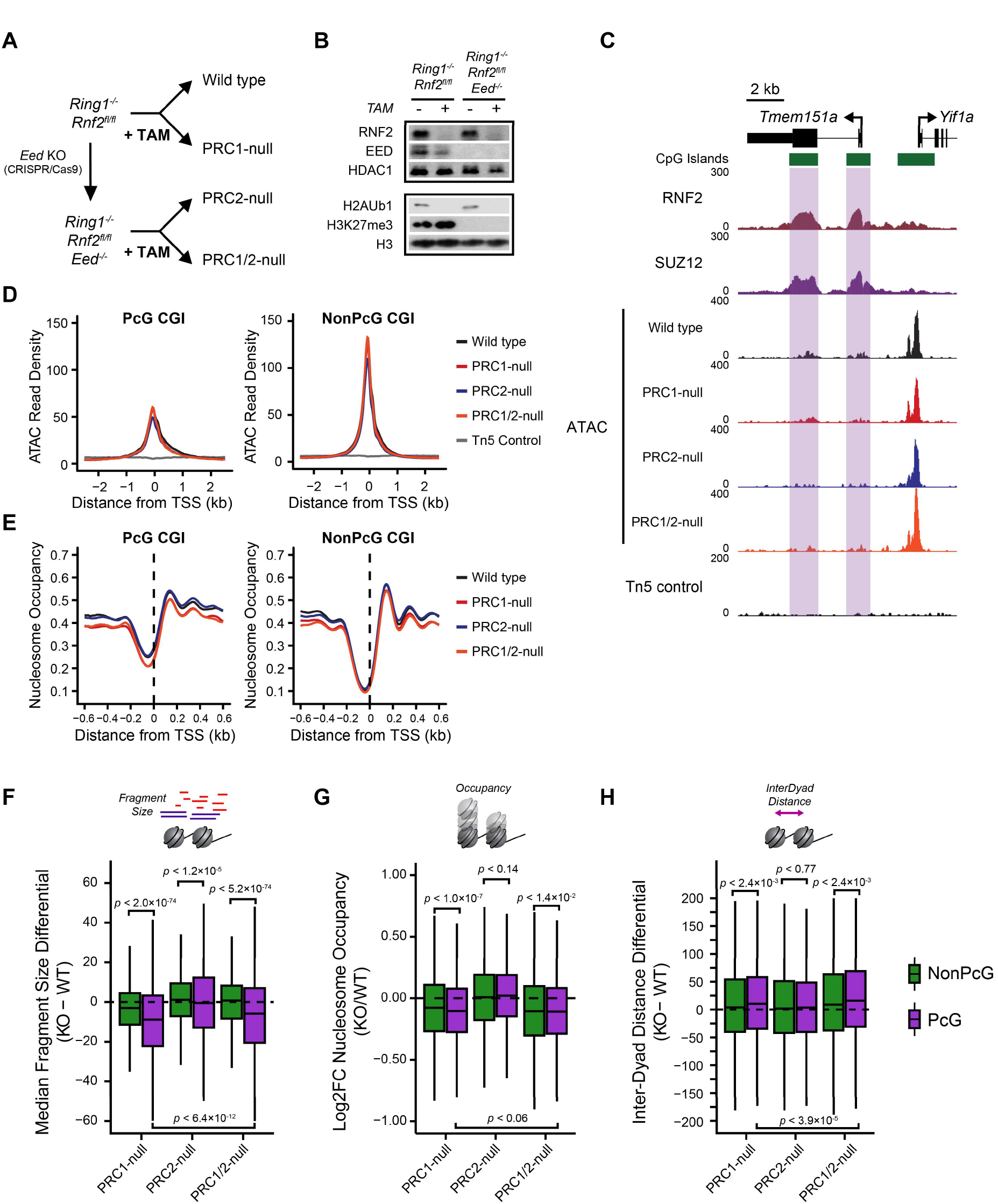
PRC1 and PRC2 do not function redundantly to shape the chromatin landscape at PcG-occupied gene promoters. A) A schematic detailing the strategy to ablate PRC1 and/or PRC2 in mouse ESCs. B) A Western blot analysis for RNF2, EED, H3K27me3 and H2AK119ub1 for *Ring1*^-/-^;*Rnf2^fl/fl^* and *Ring1*^-/-^;*Rnf2^fl/fl^*;*Eed*^-/-^ ESCs with or without tamoxifen (TAM) treatment after 72 h. C) A genome screenshot for *Ring1*^-/-^;*Rnf2^fl/fl^* and *Ring1*^-/-^;*Rnf2^fl/fl^*;*Eed*^-/-^ ATAC-seq signal before and after TAM at PcG-occupied CGIs (highlighted in purple) and a non-PcG CGI promoter. D) A metaplot analysis for *Ring1*^-/-^;*Rnf2^fl/fl^* and *Ring1*^-/-^;*Rnf2^fl/fl^*;*Eed*^-/-^ ATAC-seq before and after TAM treatment at PcG-occupied (*n* = 4020) or non-PcG CGI promoters (n = 10251), centred on TSS. E) A metaplot analysis for *Ring1*^-/-^;*Rnf2^fl/fl^* and *Ring1*^-/-^;*Rnf2^fl/fl^*;*Eed*^-/-^ NucleoATAC-derived nucleosome occupancy score before and after TAM treatment at PcG-occupied or non-PcG CGI promoters, centred on TSS. F) A boxplot comparing the change in median Tn5-tagmented fragment sizes for PcG and non-PcG CGI promoters between *Ring1*^-/-^;*Rnf2^fl/fl^* and *Ring1*^-/-^;*Rnf2^fl/fl^*;*Eed*^-/-^ ATAC-seq datasets before and after TAM treatment. G) A boxplot quantifying the log2 fold change (log2FC) in NucleoATAC-derived nucleosome occupancy for PcG and non-PcG CGI promoters in *Ring1*^-/-^;*Rnf2^fl/fl^* and *Ring1*^-/-^;*Rnf2^fl/fl^*;*Eed*^-/-^ ATAC-seq datasets before and after TAM treatment. H) A boxplot comparing the difference in median inter-dyad distances for PcG and non-PcG CGI promoters in *Ring1*^-/-^;*Rnf2^fl/fl^* and *Ring1*^-/-^;*Rnf2^fl/fl^*;*Eed*^-/-^ ATAC-seq datasets before and after TAM treatment.

### Remodelling of the PRC1-dependent nucleosome landscape is linked to RNA polymerase II activity

PcG complexes are required to maintain a transcriptionally repressive chromatin environment at developmentally-regulated gene promoters. Our characterisation of the nucleosome landscape revealed that PRC1 plays a unique role in shaping nucleosome occupancy and spacing at PcG-occupied promoters and that in the absence of PRC1 this PcG-associated nucleosome landscape reverted to an arrangement reminiscent of more transcribed non-PcG associated promoters (Figure 4). We therefore hypothesised that this altered nucleosome landscape may manifest not simply from the absence of PRC1 but instead as a result of activation of genes normally occupied by PcG complexes, potentially as a direct consequence of increased RNA polymerase II activity (Kireeva et al. 2002; Gilchrist et al. 2010; Kulaeva et al. 2010; Fenouil et al. 2012; Liang et al. 2017). This was based on our analysis that revealed correlations between gene expression, nucleosome landscape and chromatin accessibility of CGI promoters (Supplemental Figs. S1A and S3A). To test this hypothesis, we first considered whether RNA polymerase II-dependent gene transcription contributes to the nucleosome landscape or accessibility of gene promoters in a manner opposing that of PRC1. We used the chemical inhibitor triptolide to acutely inhibit RNA polymerase II initiation and occupancy prior to performing ATAC-seq (Figure 5A-D, Supplemental Fig. S7A). This resulted in significant increases in nucleosome occupancy (Figure 5B-C) and decreased distances between nucleosomal dyads in the triptolide-treated cells compared to their untreated control (Figure 5D), with the nucleosome landscape of PcG-free promoters in triptolide-treated ESCs now more closely resembling PcG-bound sites in untreated cells. This was consistent with RNA polymerase II countering the activity of PRC1 at gene promoters. However, it also suggested that the changes we observed at gene promoters in the PRC1-deficient cells might be linked to their transcriptional reactivation and not simply removal of the PRC1 complex. To examine this possibility, we performed nuclear RNA-seq in the *Ring1*^-/-^;*Rnf2^fl/fl^* and *Ring1*^-/-^;*Rnf2^fl/f^*;*Eed*^-/-^ ESCs before and after tamoxifen treatment to identify gene promoters that were activated following removal of PRC1 and/or PRC2 and directly compared these effects to alterations in the nucleosome landscape. Differential gene expression analysis identified 11.2, 14.1 and 21.2 % of all CGI promoters, and 35.6, 36.1 and 59.2 % of PcG target genes, with significant increases in gene expression after loss of PRC1, PRC2 or PRC1/2 respectively, with a high degree of overlap between cell lines and treatments (Figure 5E-F). We then compared the accessibility and nucleosome landscape at the promoters of activated genes with those whose expression was unaffected by the loss of PcG complexes. Consistent with our previous analysis, there were very few significant changes in ATAC-seq signal and these did not correlate with altered transcriptional activity at gene promoters in any of the PRC-null cell lines or triptolide-treated cells (Figure 5G, Supplemental Fig. S7B-E), demonstrating that promoter chromatin accessibility is not dependent on the transcriptional state and must be established by other mechanisms. Intriguingly, when we examined the nucleosome occupancy and spacing at PcG-bound promoters activated in the absence of PRC1 there were larger decreases in nucleosome occupancy and increased distances between nucleosome dyads compared to PcG-occupied promoters whose expression levels remained unchanged (Figure 5H-K). Although this was consistent with transcriptional changes potentially shaping the nucleosome landscape instead of a direct contribution from PRC1, we then examined PcG-bound promoters with increased activity in the PRC2-null cells. We reasoned that if RNA polymerase II and not PRC1 was responsible for the changes in the nucleosome landscape, one would expect to see comparable changes in the nucleosome landscape at upregulated gene promoters in the PRC2-null cells. However, this was not the case, as PcG target genes activated in the PRC2-null cells showed negligible or very minor differences in their nucleosome occupancy or spacing at their promoters compared to promoters with unaltered activity (Figure 5I-K). This therefore demonstrates that although RNA polymerase II activity can influence the nucleosome landscape, it is not sufficient to explain the changes that we observed at reactivated genes in the PRC1-null cells. This suggests that even in the presence of elevated transcriptional activity in PRC2-null cells, PRC1 may restrain the nucleosome landscape at these sites, potentially through disrupting RNA polymerase II-dependent chromatin remodelling. This represents a new distinction between how PRC1 and PRC2 function to shape chromatin organisation at PcG chromatin domains.

**Figure 5.**
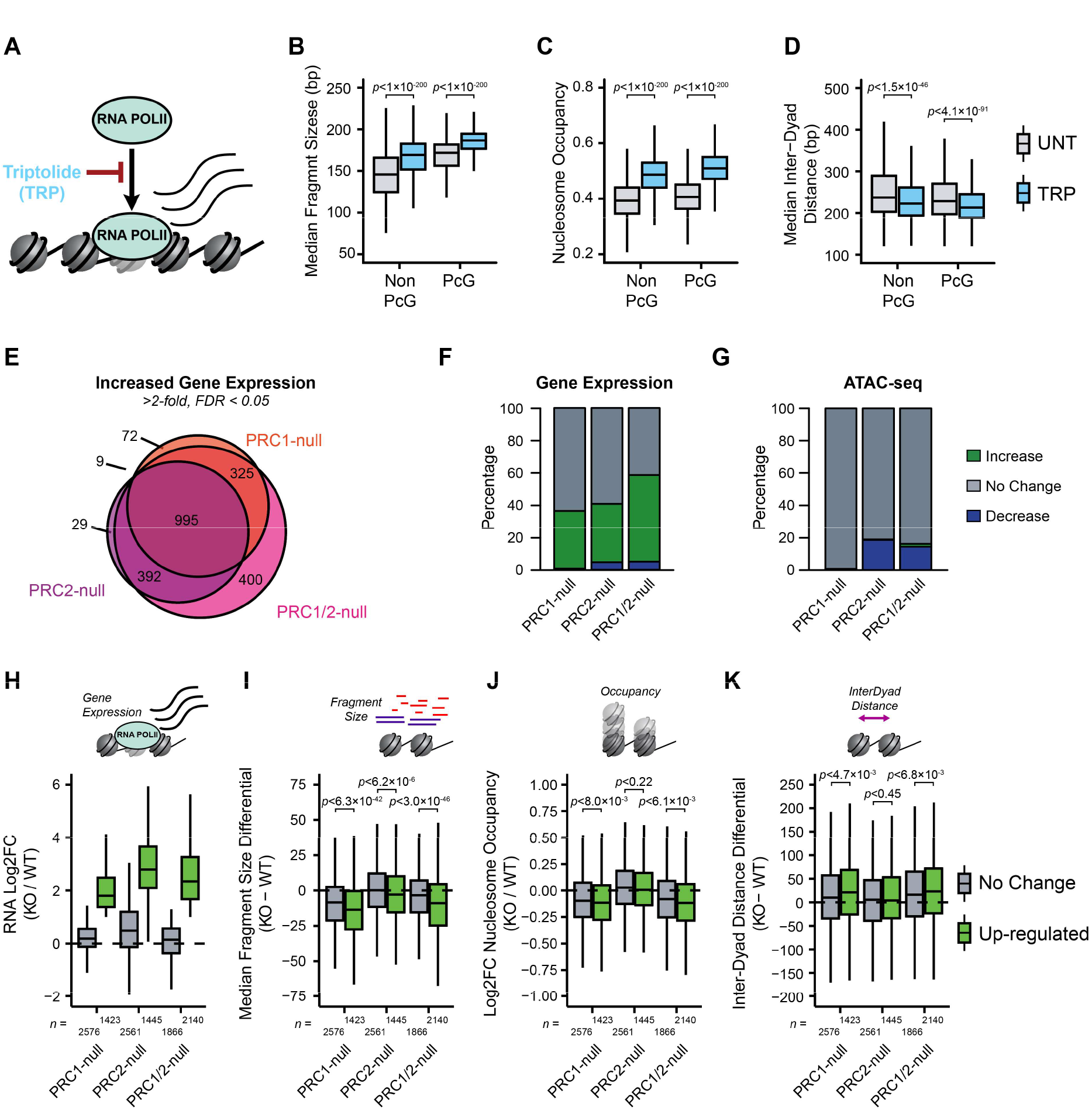
The PRC1-dependent nucleosome landscape is linked to, but not explained by, RNA polymerase II activity. A) A schematic depicting the inhibition of RNA polymerase II (RNA POLII) occupancy using triptolide (TRP). B) A boxplot comparing the median Tn5-tagmented fragment sizes for PcG and non-PcG CGI promoters before and after TRP treatment. C) A boxplot quantifying the NucleoATAC-derived nucleosome occupancy score for PcG and non-PcG CGI promoters before and after TRP treatment. D) A boxplot comparing the median inter-dyad distances for PcG and non-PcG CGI promoters before and after TRP treatment. E) A Venn diagram for PcG-occupied CGI promoters with significant increases in gene expression in *Ring1*^-/-^;*Rnf2^fl/fl^* and *Ring1*^-/-^;*Rnf2^fl/fl^*;*Eed*^-/-^ ATAC-seq before and after tamoxifen (TAM), corresponding to PRC1-null, PRC2-null and PRC1/2-null (FDR < 0.05; fold change > 2). F) A barplot depicting the number of significant RNA-seq expression changes for PcG-occupied CGI promoters. G) Same as in (F), only for ATAC-seq. Changes in ATAC-seq were calculated using the CGI promoter interval. H) A boxplot comparing the change in RNA-seq log2 fold change (log2FC) for PcG-occupied CGI promoters with (Up-regulated) or without (No Change) an increase in gene expression for each cell line and treatment. I) A boxplot comparing the change in median Tn5-tagmented fragment sizes for PcG-occupied CGI promoters with (Up-regulated) or without (No Change) an increase in gene expression for each cell line and treatment. J) A boxplot quantifying the log2 fold change (log2FC) in NucleoATAC-derived nucleosome occupancy for PcG-occupied CGI promoters with (Up-regulated) or without (No Change) an increase in gene expression for each cell line and treatment. K) A boxplot comparing the difference in median inter-dyad distances for PcG-occupied CGI promoters with (Up-regulated) or without (No Change) an increase in gene expression for each cell line and treatment.

## DISCUSSION

It has been proposed that PcG complexes establish and maintain a transcriptionally repressive chromatin state at gene promoters. *In vitro* biochemical experiments have demonstrated that PcG complexes can compact chromatin (Francis et al. 2004; Trojer et al. 2007; Grau et al. 2011; Trojer et al. 2011) and PcG-occupied gene promoters display reduced accessibility *in vivo* (Zink and Paro 1995; McCall and Bender 1996; Boivin and Dura 1998; Fitzgerald and Bender 2001; Bell et al. 2010; Calabrese et al. 2012; Beck et al. 2014; Deaton et al. 2016) (Figure 1). Here we discover that PRC1, but not PRC2, is required to maintain a chromatin landscape that is characterised by elevated nucleosome occupancy and more closely spaced nucleosomes (Figures 2-4). Unexpectedly, the ability of PRC1 to influence the local nucleosome landscape was not required to maintain the less accessible chromatin state characteristic of PcG-occupied promoters, demonstrating that chromatin compaction of reconstituted nucleosomes *in vitro* by PcG complexes cannot explain the limited accessibility at PcG target sites *in vivo*. Therefore, measurement of the accessibility and other features of the chromatin landscape at PcG targets appear not to be directly coupled, consistent with previous reports in other experimental systems (Mieczkowski et al. 2016; Mueller et al. 2017). Furthermore, we reveal that the nucleosome landscape associated with loss of PRC1 is linked to reactivation of the associated gene and may potentially reflect altered RNA polymerase II activity in the absence of PRC1 (Figure 5). Importantly, the relationship between transcriptional activation and altered nucleosome landscape was not observed in PRC2-null cells, highlighting that the transcriptionally repressive function of PRC1, but not PRC2, is linked to increased nucleosome occupancy and closer packing of nucleosomes. Together our new observations have broad implications for understanding PcG-dependent gene repression and the relationship between chromatin accessibility, the nucleosome landscape and transcriptional activity at gene promoters.

Some of the earliest studies examining chromatin at PcG-occupied sites reported reduced accessibility compared to regulatory sites lacking PcG proteins. It has been proposed that PcG protein occupancy on chromatin may define this less accessible state (McCall and Bender 1996; Fitzgerald and Bender 2001; Bell et al. 2010; Calabrese et al. 2012; Beck et al. 2014; Deaton et al. 2016). However, this possibility has not been systematically examined at the genome-scale *in vivo*. Here we directly test whether PcG proteins define accessibility at PcG-occupied gene promoters and unexpectedly find no causal relationship between the occupancy of PcG proteins and chromatin accessibility, a conclusion that was also recently reported in an independent study (Hodges et al. 2018). Therefore other activities must define the lack of accessibility at PcG genes. One possible explanation for this reduced accessibility could be that PcG target genes in ESCs have an underrepresentation of tissue-specific transcription factor binding sites which are often implicated in the recruitment of chromatin remodelling complexes such as BAF (SWI/SNF) (Ku et al. 2008; Mendenhall et al. 2010; Hu et al. 2011; Guertin and Lis 2013; Marathe et al. 2013; King and Klose 2017). In agreement with this possibility, it was recently shown that synthetic recruitment of BAF to a PcG-occupied gene promoter resulted in eviction of PcG proteins and increased accessibility (Kadoch et al. 2017). Given that we and others have shown that loss of PcG proteins is not sufficient to cause increases in chromatin accessibility at PcG target genes (Hodges et al. 2018), this increase in accessibility was presumably dependent on the chromatin remodelling activity of the BAF complex and suggests that limited activity of BAF or possibly other chromatin remodelling complexes may explain the low accessibility of PcG-occupied chromatin. Another feature of PcG-occupied promoters is their low levels of histone acetylation compared to other gene promoters. Histone acetylation is associated with elevated chromatin accessibility (Rincon-Arano et al. 2012; Lennartsson et al. 2015; Frank et al. 2016). At PcG targets, the removal of acetylation from lysine residues in histone tails would reinstate their positive charge and allow them to more stably interact with DNA and possibly limit accessibility (Norton et al. 1989; Shogren-Knaak et al. 2006). This could be mediated by the nucleosome remodelling and deacetylase (NuRD) complex which co-occupies many PcG target sites (Yildirim et al. 2011; Reynolds et al. 2012) and has been demonstrated to limit chromatin accessibility at regulatory elements (Ramírez et al. 2012; de Dieuleveult et al. 2016). Ultimately, it remains unclear what defines reduced chromatin accessibility at PcG targets, and what, if any, role this plays in regulating gene expression at these sites.

Our analysis of the nucleosome landscape at PcG target sites revealed a role for PRC1 in regulating nucleosome occupancy and spacing that is distinct from chromatin accessibility. One would have predicted that the reduced occupancy of nucleosomes following removal of PRC1 would yield an increase in chromatin accessibility, however this is not evident in our analysis. The precise molecular explanation for this discord remains unclear. One possibility is that alternative proteins engage with sites vacated by PcG proteins competing with nucleosomes for occupancy, but not affecting overall accessibility measurements. Alternatively, the effects we observe on the nucleosome landscape in PRC1-null cells are very subtle and may not lead to a profound enough perturbation of local chromatin structure to manifest in overall increases in chromatin accessibility. Nevertheless, a lack of concordance between the measurement of accessibility and nucleosome features has been reported previously (Mieczkowski et al. 2016; Mueller et al. 2017), indicating that the relationship between these measurements is not always simple to rationalise. Clearly in future work it will be important to understand in more detail how the nucleosome landscape of gene promoters is related to measurements of chromatin accessibility, particularly in the context of PcG-bound sites.

Here we have disrupted PRC1 by removing the core scaffolding proteins RING1/RNF2 which are also the E3 ubiquitin ligases required for deposition of H2AK119ub1. PRC1 has been proposed to function through E3 ligase-dependent and -independent activities (Endoh et al. 2012; Blackledge et al. 2014; Cooper et al. 2014; Illingworth et al. 2015; Pengelly et al. 2015; Rose et al. 2016) and its ability to compact chromatin *in vitro* is thought to be independent of its ubiquitin ligase activity (Francis et al. 2004; Margueron et al. 2008). It will be interesting to determine if these E3 ligase-independent activities characterised *in vitro* contribute to PRC1’s effect on the nucleosome landscape *in vivo* by examining PcG-occupied chromatin in situations where the E3 ligase activity of RING1/RNF2 has been eliminated (Endoh et al. 2012; Illingworth et al. 2015). However, if the catalytic activity of PRC1 is not responsible for shaping the nucleosome landscape, how could this be achieved? Two PRC1 components linked to chromatin compaction and the inhibition of chromatin remodelling *in vitro*, BMI1 (also known as PCGF4) and CBX2, contain highly basic and disordered protein domains that are conserved across different PcG components in different species (Grau et al. 2011; Beh et al. 2012). Increasing the acidity of this domain in CBX2 disrupted its ability to inhibit chromatin remodelling (Grau et al. 2011), suggesting that the presence of these basic and highly charged domains might also be important for the *in vivo* regulation of nucleosome occupancy and spacing. However, both CBX2 and BMI1 are expressed at low levels in ESCs and form only a small minority of PRC1 complexes (Kloet et al. 2016), so it is unclear what their contribution towards the PcG-dependent nucleosome landscape could be in this cell type. Finally, several studies support the possibility that PRC1 might interfere directly with RNA polymerase II occupancy or activity (Stock et al. 2007; Brookes et al. 2012; Lehmann et al. 2012). In agreement with these findings, following deletion of PRC1 we observed that a subset of promoters is susceptible to transcriptional activation and these then acquire a nucleosome landscape consistent with elevated RNA polymerase II activity. Alterations in the nucleosome landscape following PRC1 removal are therefore likely driven by processes linked to transcription. However, elevated transcription *per se* is not necessarily sufficient to drive these outcomes, as some PcG target genes display elevated expression following removal of PRC2, yet nevertheless, retain a PRC1-dependent nucleosome landscape. Investigating the detailed mechanisms that define the nucleosome landscape at PcG target genes and how this is related to gene transcription will be an interesting area for future work and will be fundamental to understanding how PcG complexes repress gene transcription.

In conclusion, we have discovered that PRC1 can influence the nucleosome landscape at PcG target genes in a manner that does not contribute to reduced chromatin accessibility. This indicates that PRC1-dependent chromatin compaction observed *in vitro* does not explain the reduced accessibility at PcG target sites *in vivo* and reveals a new and previously unappreciated complexity in the relationship between PcG complexes, the nucleosome landscape and gene repression.

## METHODS

### Cell culture and lines

Mouse embryonic stem cell (ESC) lines were grown on gelatin-coated plates in DMEM supplemented with 15 % FBS, 10 ng/mL leukemia-inhibitory factor, penicillin/streptomycin, β-mercaptoethanol, L-glutamine and non-essential amino acids. *Ring1*^-/-^;*Rnf2^fl/fl^* ESCs (Endoh et al. 2008) were adapted to grow under feeder-free culture conditions and were treated with 800 nM 4-hydroxytamoxifen (TAM) for 72 h to ablate RNF2 levels. EED conditional knockout ESCs that express a doxycycline-sensitive *Eed4* transgene (*Eed4*^TG^) in an *Eed*^-/-^ background were treated with 1 μg/mL doxycycline (DOX) for 14 days to disrupt PRC2 complex and function, as previously described (Ura et al. 2008; Tavares et al. 2012). SV40-immortalised *Ring1*^-/-^;*Rnf2^fl/fl^* mouse embryonic fibroblasts (MEFs) (Endoh et al. 2012; Jullien et al. 2017) were grown in DMEM supplemented with 7% FBS and penicillin/streptomycin and maintained in culture for less than 10 passages. *Ring1*^-/-^;*Rnf2^fl/fl^* MEFs were treated with 800 nM TAM for 96 h to ablate RNF2 levels. Loss of protein expression and Polycomb complex activity was verified by Western blotting using the following antibodies: RNF2 (Blackledge et al. 2014), SMARCA4 (abcam, ab110641), EED (Millipore, #09-774), HDAC1 (abcam, ab109411), H2AK119ub1 (Cell Signalling Technology (CST), #8240), H3K27me3 (Diagenode, pAb-069-050 and (Rose et al. 2016)), H3 (Farcas et al. 2012), H2A (CST, #3636), H4 (CST, #2935). All cell lines were confirmed to be mycoplasma-free.

### Generation of Polycomb double-knockout ESCs

To delete EED in the *Ring1*^-/-^;*Rnf2^fl/fl^* ESCs, CRISPR/Cas9 guides were designed flanking exons 2 to 5 of *Eed* (Guide 1: 5’ CACCGACAATCAGTGCTCTTACTCG 3’; Guide 2: 5’ CACCGAAACAGTAAGAGTCGAGTCG 3’) to induce a frameshift in all four EED translation products. The *Eed* sgRNAs were cloned into pSpCas9(BB)-2A-Puro (plasmid 48139; Addgene, Cambridge, MA) using a previously described protocol (Ran et al. 2013). Lipofectamine 3000 (Life Technologies) was used to transfect Cas9-sgRNA plasmids into *Ring1*^-/-^;*Rnf2^fl/fl^* ESCs and transfected cells were treated with 1 μg/mL puromycin for 48 hr. After 10 days, individual colonies were isolated, expanded and genomic DNA was screened by PCR for deletion of *Eed* exons 2 to 5 (FWD: 5’ AGCAGGCAGATACCAGAGTG 3’; REV 5’ ATGTCAGCACGTCCCAACTA 3’). Putative *Eed*^-/-^ clones were confirmed by Western blotting. *Ring1*^-/-^;*Rnf2^fl/fl^*;*Eed*^-/-^ cells were treated with 800 nM TAM for 72 h to ablate RNF2 expression.

### Inhibition of RNA polymerase II

To inhibit RNA polymerase II activity, E14 ESCs were pre-plated at 2.5×10^6^ cells/10 cm plate and allowed to grow for 24 h prior to treatment with 500 nM triptolide (TRP) for 50 min, as previously described (Jonkers et al. 2014). To limit re-activation of RNA polymerase II, cells were immediately washed with ice-cold PBS and harvested by cell scraping prior to nuclei isolation for RNA and ATAC analysis. To validate TRP treatment, real-time reverse transcriptase PCR was performed using intronic (pre-mRNA) primer sequences and normalised to the RNA polymerase III-transcribed U6 snRNA gene using the ΔΔCt method.

### ATAC-seq sample preparation and sequencing

Chromatin accessibility was assayed using an adaptation of the assay for transposase accessible-chromatin (ATAC)-seq (Buenrostro et al. 2013), as previously described (King and Klose 2017). Briefly, nuclei were isolated in 1 mL HS Lysis buffer (50 mM KCl, 10 mM MgSO4.7H20, 5 mM HEPES, 0.05 % NP40 (IGEPAL CA630)), 1 mM PMSF, 3 mM DTT) for 1 min at room temperature and washed three times with ice-cold RSB buffer (10 mM NaCl, 10 mM Tris (pH 7.4), 3 mM MgCl2). 5×10^4^ nuclei were counted and resuspended in 1X Tn5 reaction buffer (10 mM TAPS, 5 mM MgCl2, 10 % dimethylformamide) with 2 μl of Tn5 transposase (25 μM) made in house according to the previously described protocol (Picelli et al. 2014). Reactions were incubated for 30 min at 37°C, before isolation and purification of tagmented DNA using QiaQuick MinElute columns (Qiagen). ATAC-seq libraries were prepared by PCR amplification using single index (i7) Illumina barcodes previously described (Buenrostro et al. 2013) and the NEBNext^®^ High-Fidelity 2X PCR Master Mix with 8-10 cycles. Libraries were quantified by qPCR using SensiMix SYBR (Bioline) and KAPA Library Quantification DNA standards (KAPA Biosystems), and sequenced on Illumina NextSeq500 using 80 bp paired-end reads in biological duplicate (*Eed*^-/-^;*Eed4.TG^DOX^*), triplicate (TRP treatment and *Ring1*^-/-^;*Rnf2^fl/fl^* MEFs) or quadruplicate (*Ring1*^-/-^;*Rnf2^fl/fl^* and *Ring1*^-/-^;*Rnf2^fl/fl^*;*Eed*^-/-^).

### MNase-seq sample preparation and sequencing

For micrococcal nuclease (MNase)-seq experiments, we used an adaptation of a native ChIP protocol described previously (Rose et al. 2016). Briefly, nuclei were isolated from 5 × 10^7^ *Ring1*^-/-^;*Rnf2^fl/fl^* mouse embryonic stem cells with or without TAM-treatment with RSB buffer (10 mM Tris-HCl (pH 8), 10 mM NaCl, 3 mM MgCl2) supplemented with 0.1 % NP-40 and 5 mM N-ethylmaleimide. This was followed by digestion for 5 min at 37°C with 16 U MNase (Fermentas, Waltham, MA) in 1 ml RSB supplemented with 0.25 M sucrose, 3 mM CaCl2 and 10 mM N-ethylmaleimide. After digestions were stopped with 4 mM EDTA, nuclei were pelleted by centrifugation at 1500 x *g* and the soluble S1 fraction collected. Pelleted nuclei were then resuspended in 300 μl nucleosome release buffer (10 mM Tris-HCl (pH 7.5), 10 mM NaCl, 0.2 mM EDTA, 10 mM N-ethylmaleimide), incubated at 4°C for 1 hr with gentle rotation, and then gently passed through a 27G syringe needle five times. After the insoluble material was pelleted by centrifugation at 1500 x g, the soluble S2 fraction was collected and combined with the S1 fraction. To prepare material for constructing sequencing libraries, DNA was purified from chromatin corresponding to 5 × 10^6^ cells using ChIP DNA Clean and Concentrator kit (Zymo, Irvine, CA). The efficiency of MNase digestion was assessed by DNA electrophoresis (1.5% agarose gel). MNase-seq libraries were prepared from 500 ng of DNA using the NEBNext Ultra II DNA Library Prep Kit (NEB) according to the manufacturer’s protocol. DNA fragment size selection step was included to enrich for mono-nucleosome size fragments in the final libraries. Libraries were quantified as for ATAC-seq libraries and were sequenced on Illumina NextSeq500 using 80 bp paired-end reads in biological triplicate.

### Nuclear RNA-seq sample preparation and sequencing

To purify nuclear RNA, nuclei were isolated as described for ATAC-seq prior to resuspension in TriZOL reagent (ThermoScientific) and RNA extraction according to the manufacturer’s protocol. RNA was treated with the TURBO DNA-free Kit (ThermoScientific) and rRNA was depleted using the NEBNext rRNA Depletion kit (NEB). RNA-seq libraries were prepared using the NEBNext Ultra Directional RNA-seq kit (NEB) and libraries were sequenced on the Illumina NextSeq500 with 80 bp paired-end reads in biological quadruplicate.

### Sequencing data alignment, processing and normalisation

For ATAC-seq, DNase-seq, FAIRE-seq, MNase-seq, ChIP-seq and BioCAP-seq datasets paired-end reads were aligned to the mouse mm10 genome using bowtie2 (Langmead and Salzberg 2012) with the “--no-mixed” and “--no-discordant” options, while single-end libraries were aligned using default bowtie2 settings. Non-uniquely mapping reads and reads mapping to a custom blacklist of artificially high regions of the genome were discarded. For RNA-seq, reads were initially aligned using bowtie2 against the rRNA genomic sequence (GenBank: BK000964.3) to quantify and filter out rRNA fragments, prior to alignment against the mm10 genome using the STAR RNA-seq aligner (Dobin et al. 2012). PCR duplicates were removed using SAMtools (Li et al. 2009). Biological replicates were randomly downsampled to contain the same number of reads for each individual replicate, and merged to create a representative genome track using DANPOS2 (Chen et al. 2013) for ATAC-seq and MNase-seq samples, MACS2 (Zhang et al. 2008) for ChIP-seq, FAIRE-seq and BioCAP-seq or genomeCoverageBed (Quinlan 2014) for RNA-seq. Genome coverage tracks were visualised using the UCSC Genome Browser (Kent et al. 2002).

### Differential accessibility and gene expression analysis

Significant changes in ATAC-seq datasets were identified using the DiffBind package (Stark and Brown 2011), while for RNA-seq DESeq2 (Love et al. 2014) was used with a custom-built, non-redundant mm10 gene set (Rose et al. 2016). For DiffBind analysis of ATAC-seq datasets *FDR* < 0.05 and a fold change > 1.5-fold was deemed a significant change, while for DESeq2 analysis of RNA-seq a threshold of *FDR* < 0.05 and a fold change > 2-fold was used.

### Annotation and analysis of Polycomb target sites

Non-redundant refGene TSS intervals (±500bp; *n* = 20633) were overlapped with mouse ESC RNF2 and SUZ12 peak sets previously identified from biological triplicate data with input control using MACS2 (Rose et al. 2016), and any TSS overlapping with both RNF2 and SUZ12 were considered to be *bona fide* Polycomb target TSS. Non-methylated CpG island (CGI) intervals were experimentally identified in ESCs using MACS2 peak calling of BioCAP-seq (Blackledge et al. 2012; Long et al. 2013), and only TSS within CGI intervals were used for subsequent promoter-based analyses. For MEF and mouse ENCODE tissues, CGI intervals downloaded from the UCSC Genome Browser that overlapped with the non-redundant set of TSS were annotated with bisulfite sequencing methylation profiles (Hon et al. 2013; Hu et al. 2014) lifted over to mm10 using the liftOver tool from UCSC (Hinrichs et al. 2006) and only intervals with methylation <5% were considered. For each tissue or cell line polycomb target CGI TSS were identified by overlapping with ENCODE H3K27me3 peaks (Yue et al. 2014) lifted over from mm9 to mm10 or identified using MACS2 peak calling (Han et al. 2017). To identify Polycomb-bound distal regulatory elements in mouse ESCs, ATAC peaks identified with DANPOS2 in wild type ESCs were annotated with H3K4me1 (Whyte et al. 2012) and H3K4me3 (Yue et al. 2014) to classify putative distal regulatory elements as previously described (King and Klose 2017), and peaks overlapping with both RNF2 and SUZ12 were considered Polycomb targets. Gene expression-matched promoters were identified using untreated *Ring1*^-/-^;*Rnf2^fl/fl^* ESC nuclear RNA-seq normalised expression values calculated by DESeq2. Metaplot analysis of ATAC-seq, MNase-seq, ChIP-seq or nucleosome occupancy profiles at gene promoters was performed using HOMER2 (Heinz et al. 2010). Quantitation of reads per kilobase per million (RPKM) was performed within CGI intervals at TSS using custom scripts. Data were visualised using R (v 3.2.1) and ggplot2, with scatterplots coloured by density using stat_density2d. Regression and correlation analyses were also performed in R using standard linear models and Pearson correlation respectively.

### Characterisation of nucleosome features at gene promoters

As a simple measure of nucleosome occupancy at promoters, the fragment sizes of Tn5-tagmented DNA fragments within each promoter interval were extracted from ATAC-seq .bam files and used to calculate the median fragment size per CGI promoter interval. Higher median fragment sizes correspond to higher levels of nucleosome-sized Tn5-tagmented DNA, while lower fragment sizes correspond to higher levels of nucleosome-free DNA. To complement this approach, we extracted signal corresponding to nucleosome occupancy and positional information within CGI promoters using the NucleoATAC package (Schep et al. 2015), which relies upon a model-based analysis of Tn5 tagmentation fragment size profiles to reflect the probability of nucleosome occupancy at a given loci. Importantly, both methods are independent of the total coverage of tagmented fragments (i.e. accessibility) at different loci. For MNase-seq datasets, nucleosome positions and occupancy were determined using DANPOS2 (Chen et al. 2013). In order to visualise nucleosome occupancy, we profiled the occ.bedgraph files from our NucleoATAC analysis and normalised .wig for MNase-seq tracks centred upon TSS in 1bp resolution and identified average nucleosome positions using the local maxima of the coverage. Quantification of total nucleosome occupancy per kb for CGI promoters was performed by calculating the coverage of NucleoATAC-derived .occ.bedgraph files using bedtools “coverage” tool (Quinlan 2014) or the median nucleosome summit height from DANPOS2 MNase-seq nucleosome calls per CGI. Individual nucleosome dyad centres were identified in the nucmap_combined.bed file from NucleoATAC and DANPOS2 MNase-seq nucleosome calls and were used to calculate the distance to the nearest neighbouring nucleosome dyad centre (inter-dyad distance) using the bedtools “closest” tool. Only nucleosomes within CGI intervals were included for this analysis, and the median inter-dyad distance for each CGI interval was calculated. Median nucleosome fuzziness scores per CGI were calculated from NucleoATAC-derived nucpos.bed files or DANPOS2 MNase-seq nucleosome calls.

## DATA ACCESS

The ATAC-seq, MNase-seq and RNA-seq data from this study have been submitted to the NCBI Gene Expression Omnibus (GEO; https://www.ncbi.nlm.nih.gov/geo/) under accession number GSE98403. Previously published datasets used for analysis include mouse ESC DNase-seq (GSE37074; Yue et al. 2014), FAIRE-seq (GSE49141; Thakurela et al. 2013), Tn5 digestion control (GSE87822; King and Klose 2017), RNF2 and SUZ12 ChIP-seq (GSE83135; Rose et al. 2016), BioCAP (GSE43512; Long et al. 2013), H3K4me1 ChIP-seq (GSE27844; Whyte et al. 2012) and H3K4me3 ChIP-seq (GSE49847; Yue et al. 2014), mouse tissue H3K27me3 (GSE49847; Yue et al. 2014) and whole genome bisulfite sequencing (GSE42836; Hon et al. 2013), MEF H3K27me3 (GSE91374; Han et al. 2017) and reduced representation bisulfite sequencing (GSE52741; Hu et al. 2014), and triptolide-treated GRO-seq (GSE48895; Jonkers et al. 2014).

## ACKNOWLEDGEMENTS

Work in the Klose lab is supported by the Wellcome Trust, the Lister Institute of Preventive Medicine, EMBO and the European Research Council. We would also like to thank everyone in the Klose lab for advice and support.

---

**Supplemental Figure S1.**
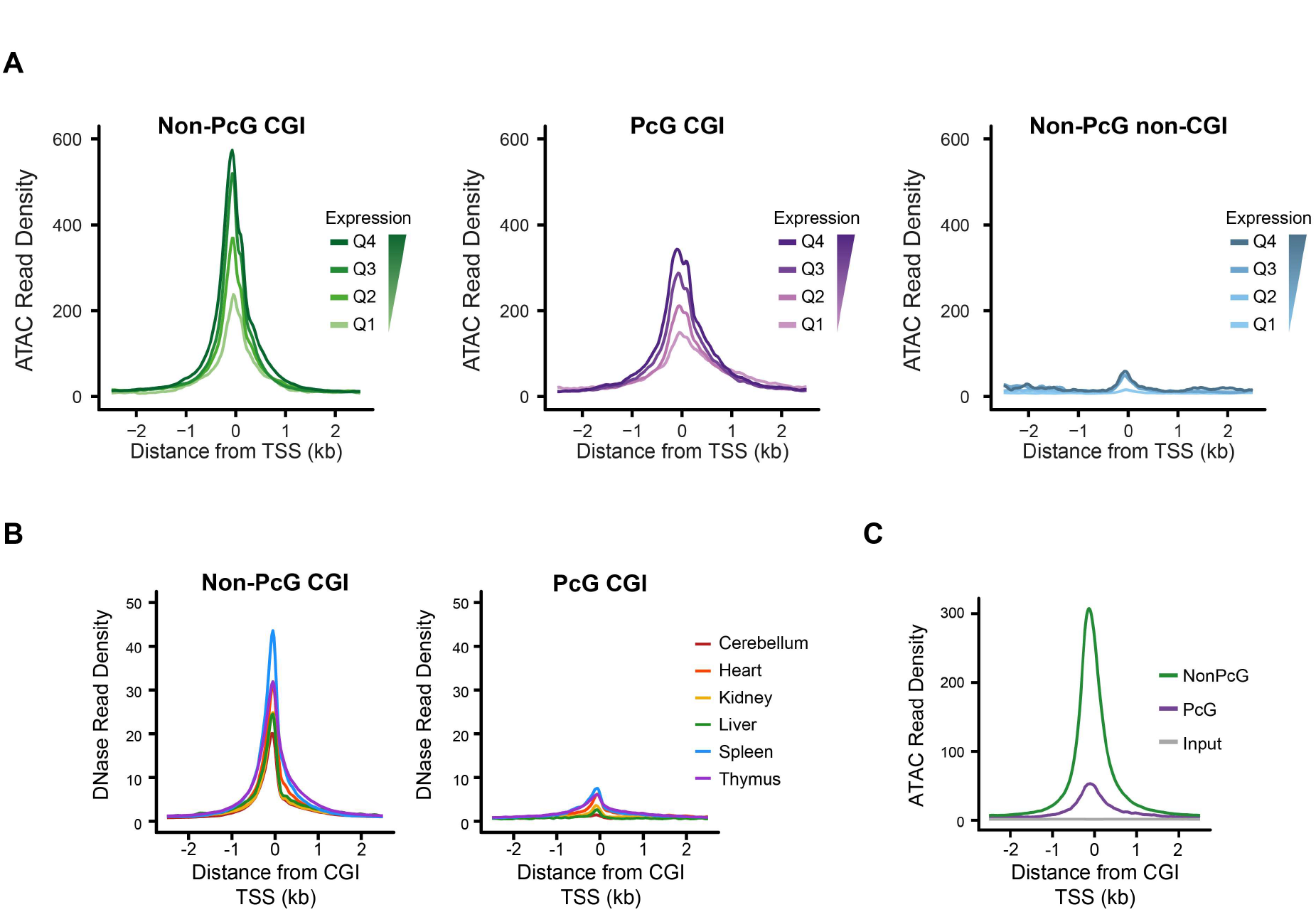
Analysis of chromatin accessibility at Polycomb and non-Polycomb CGI promoters. A) A metaplot analysis of wild type mouse ESC ATAC-seq profiles at non-PcG CGI, PcG-bound CGI and non-CGI PcG-free promoters at different gene expression quartiles (Q1 lowest -> Q4 highest), centred on the TSS. B) A metaplot analysis of ENCODE DNase-seq for different mouse tissues at H3K27me3-positive or H3K27me3-negative non-methylated CGI promoters, centred on the TSS. C) A metaplot analysis of wild type mouse embryonic fibroblast ATAC-seq profiles at H3K27me3-positive (PcG; *n* = 1438) or H3K27me3-negative CGI promoters (Non-PcG; *n* = 10118), centred on the TSS.

**Supplemental Figure S2.**
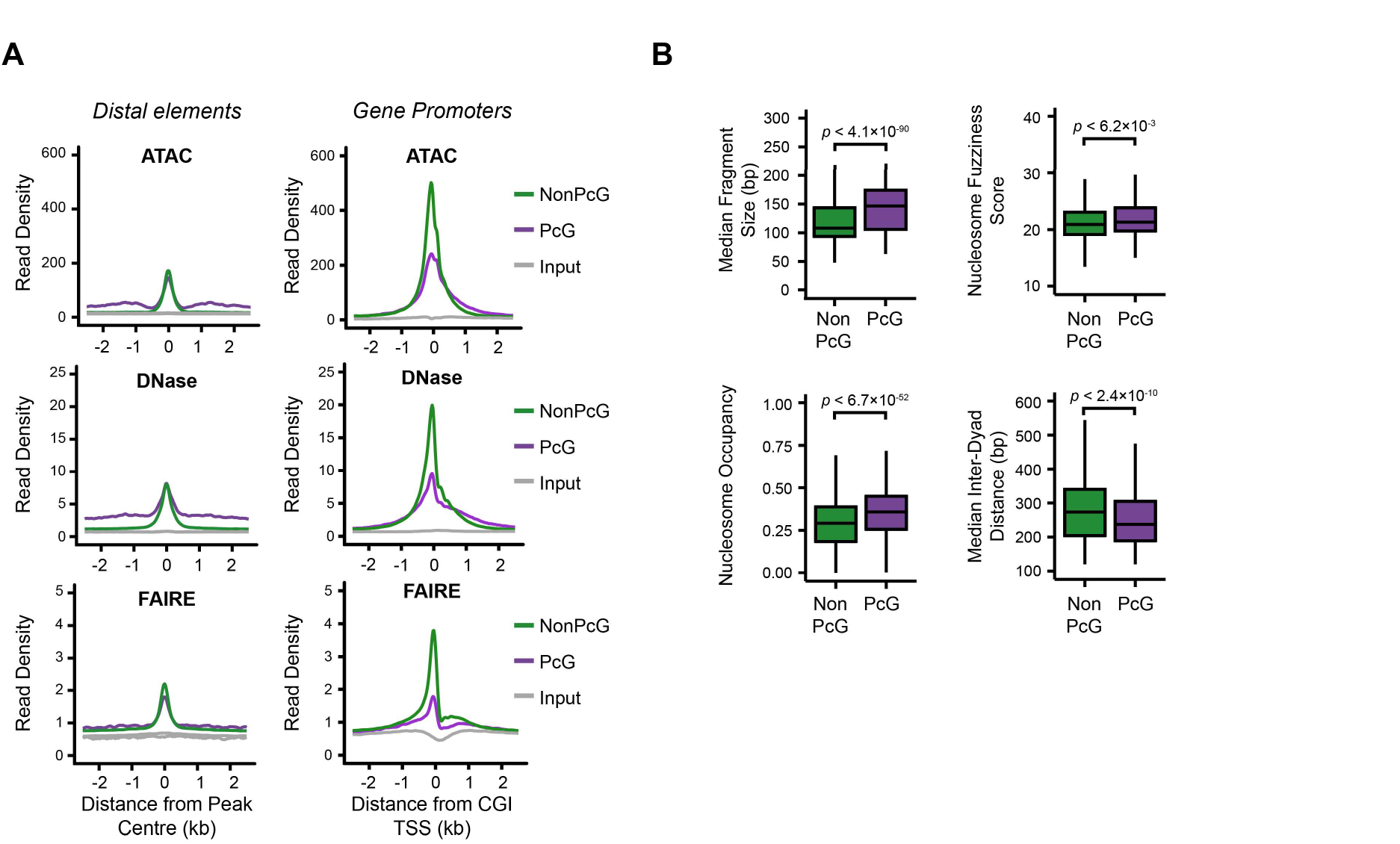
Chromatin accessibility and nucleosome landscape features at distal elements bound by Polycomb. A) A metaplot analysis of ATAC-seq, DNase-seq and FAIRE-seq signal at PcG-occupied (*n* = 1229) or PcG-free (*n* = 43134) distal elements (left) compared with CGI promoters (right; as in 1D). B) Boxplots comparing the median ATAC-seq fragment sizes, NucleoATAC-derived nucleosome occupancy signal, nucleosome fuzziness and median inter-dyad distances for PcG-occupied (*n* = 4020) or non-PcG (*n* = 10251) CGI promoters.

**Supplemental Figure S3.**
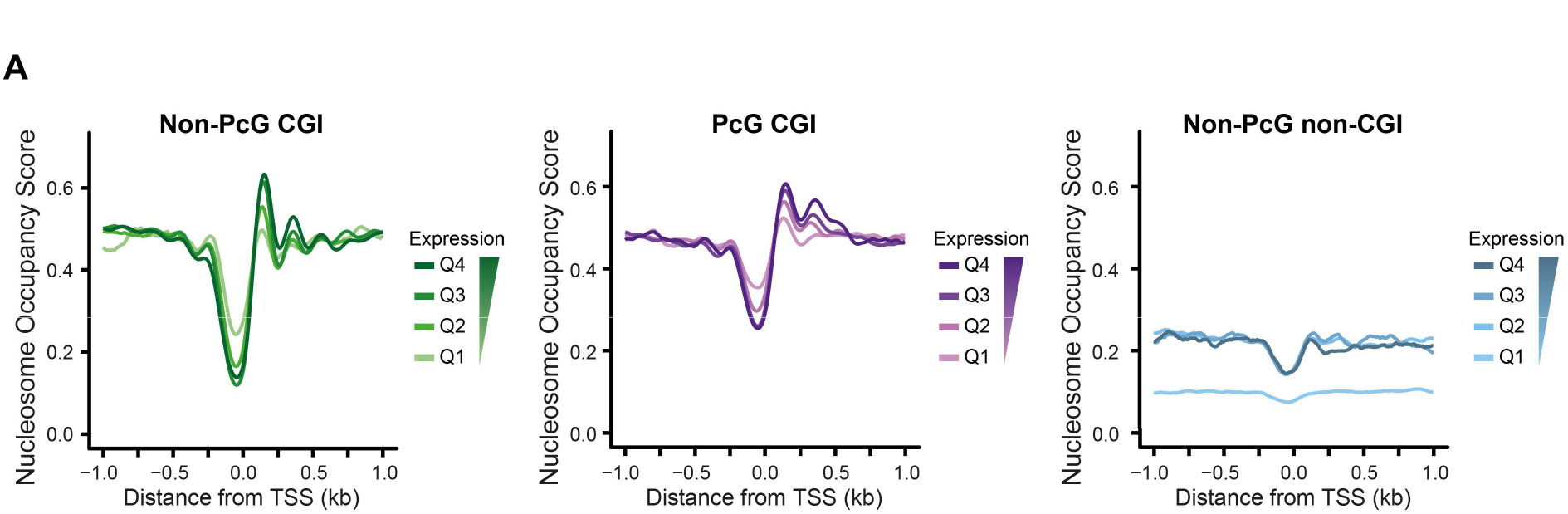
Nucleosome landscape of expression-matched PcG-bound or PcG-free promoters. A) A metaplot analysis of wild type mouse ESC NucleoATAC-derived nucleosome occupancy scores at non-PcG CGI, PcG-bound CGI and non-CGI PcG-free promoters of different gene expression quartiles (Q1 lowest -> Q4 highest).

**Supplemental Figure S4.**
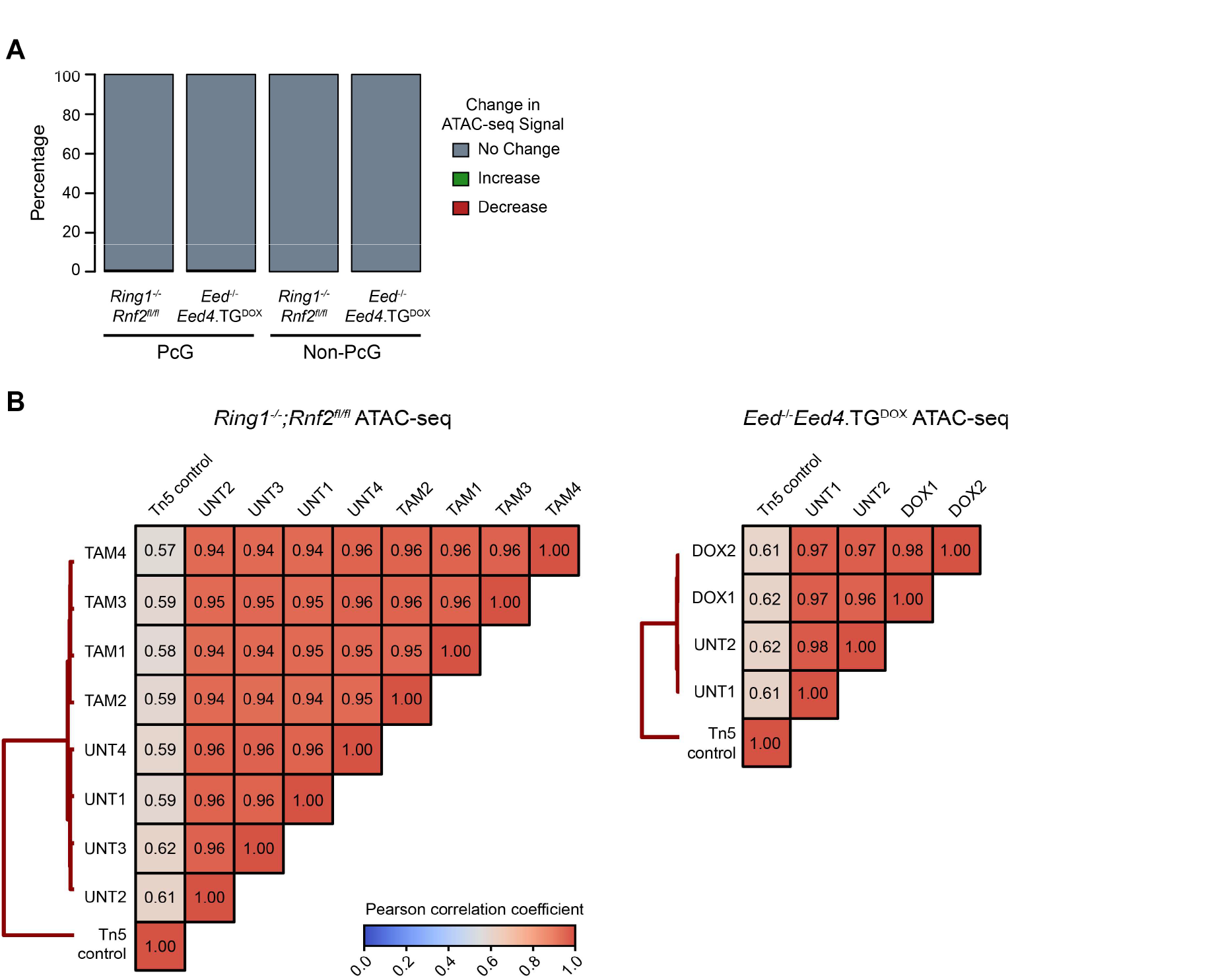
Statistical analysis and reproducibility of ATAC-seq datasets. A) A barplot depicting the number of significant changes in ATAC-seq signal at PcG or Non-PcG CGI promoters after ablation of PRC1 (*Ring1*^-/-^;*Rnf2^fl/fl^*) or PRC2 (*Eed*^-/-^;*Eed4.TG^DOX^*). B) Pearson correlation matrices for biological replicate ATAC-seq signal at wild type ESC ATAC hypersensitive sites for *Ring1*^-/-^;*Rnf2^fl/fl^* and *Eed*^-/-^;*Eed4.TG^DOX^* ATAC-seq experiments.

**Supplemental Figure S5.**
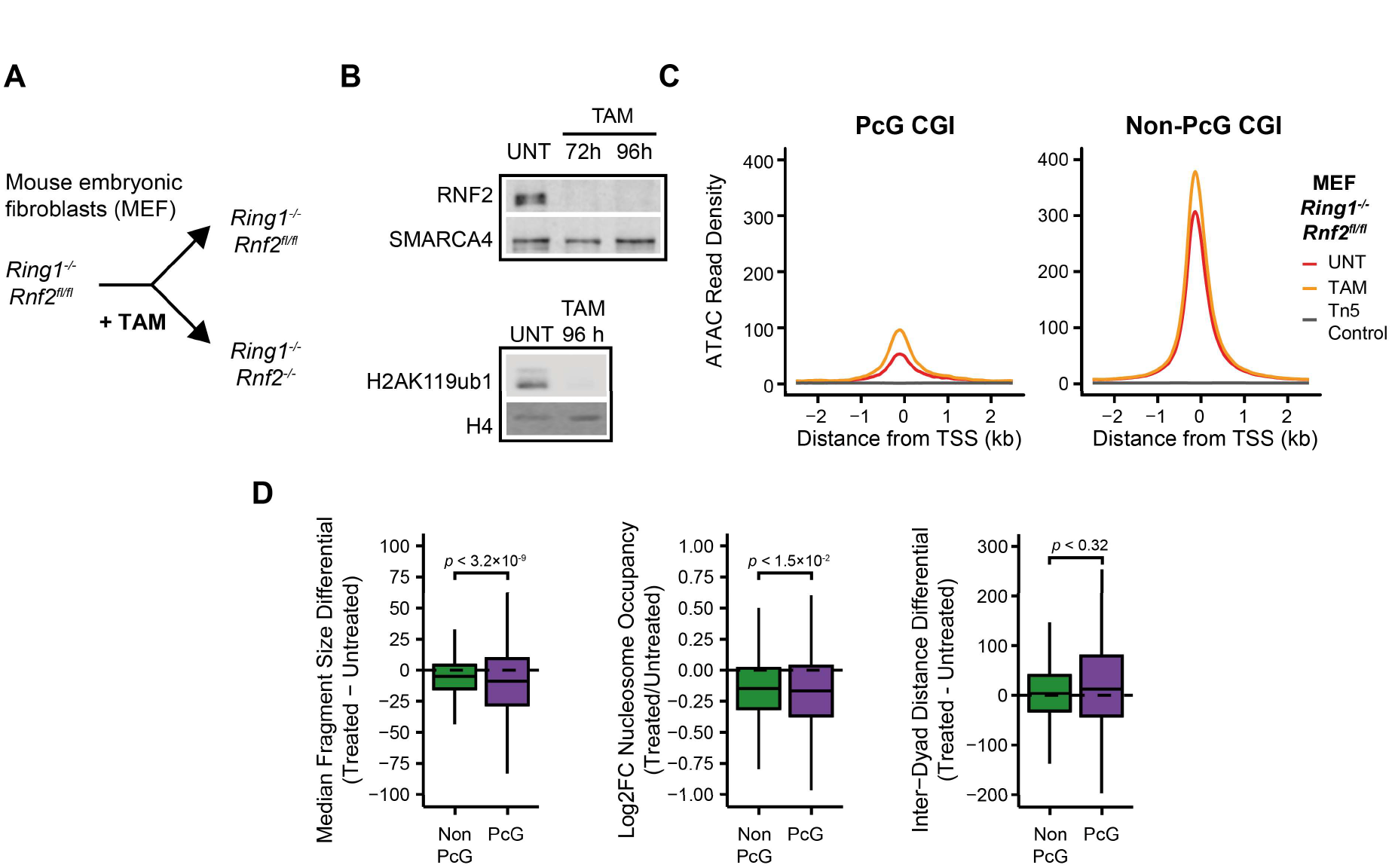
Analysis of chromatin accessibility and the nucleosome landscape in PRC1-null mouse embryonic fibroblasts. A) A schematic depicting the treatment of *Ring1*^-/-^;*Rnf2^fl/fl^* mouse embryonic fibroblasts (MEFs) with 4-hydroxytamoxifen (TAM) to generate PRC1-null MEFs. B) A Western blot analysis of untreated and TAM-treated *Ring1*^-/-^;*Rnf2^fl/fl^* MEFs for RNF2 (upper) and H2AK119ub1 (lower) at different time points. 96 hours after TAM treatment was used for all future experiments. C) A metaplot analysis for *Ring1*^-/-^;*Rnf2^fl/fl^* MEF ATAC-seq before and after tamoxifen (TAM) treatment at H3K27me3-positive (PcG; *n* = 1438) or H3K27me3-negative CGI promoters (Non-PcG; *n* = 10118). D) Quantitation of differences in median ATAC-seq fragment sizes, NucleoATAC-derived nucleosome occupancy, and inter-dyad distances for PcG and non-PcG CGI promoters in *Ring1*^-/-^;*Rnf2^fl/fl^* MEF ATAC-seq before and after TAM treatment.

**Supplemental Figure S6.**
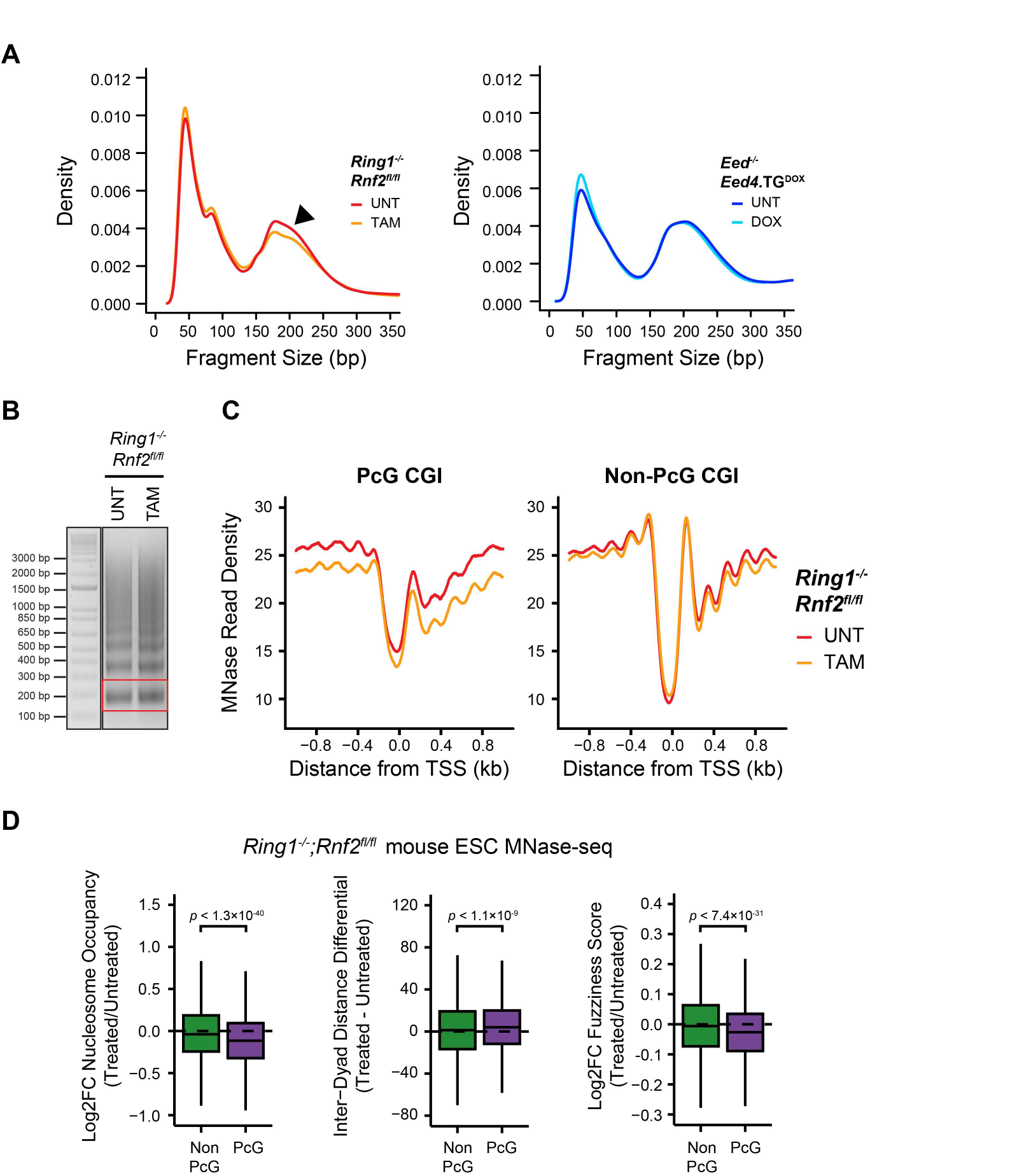
Deletion of PRC1 results in an altered nucleosome landscape. A) Frequency distribution plots for ATAC-seq fragment sizes in PcG-occupied CGI promoter intervals in *Ring1*^-/-^;*Rnf2^fl/fl^* and *Eed*^-/-^;*Eed4.TG^DOX^* ESC ATAC-seq with or without tamoxifen (TAM) or doxycycline (DOX) treatment respectively. The arrow highlights loss of mono-nucleosome-sized fragments in the *Ring1*^-/-^;*Rnf2^fl/fl^* experiment. B) An exemplar agarose gel (1.5 %) electrophoresis of MNase-digested native chromatin of a single replicate of MNase-seq library construction for *Ring1*^-/-^;*Rnf2^fl/fl^* ESCs. The red box highlights the mono-nucleosome fraction that was size-selected during library preparation. C) A metaplot analysis for *Ring1*^-/-^;*Rnf2^fl/fl^* MNase-seq before and after TAM treatment at PcG-occupied (*n* = 4020) or PcG-free (*n* = 10251) CGI promoters centred on the TSS. D) Comparison of differences in MNase-seq derived measurements of nucleosome occupancy, interdyad distance and nucleosome fuzziness for PcG and non-PcG CGI promoters in *Ring1*^-/-^;*Rnf2^fl/fl^* ESCs before and after TAM treatment.

**Supplemental Figure S7.**
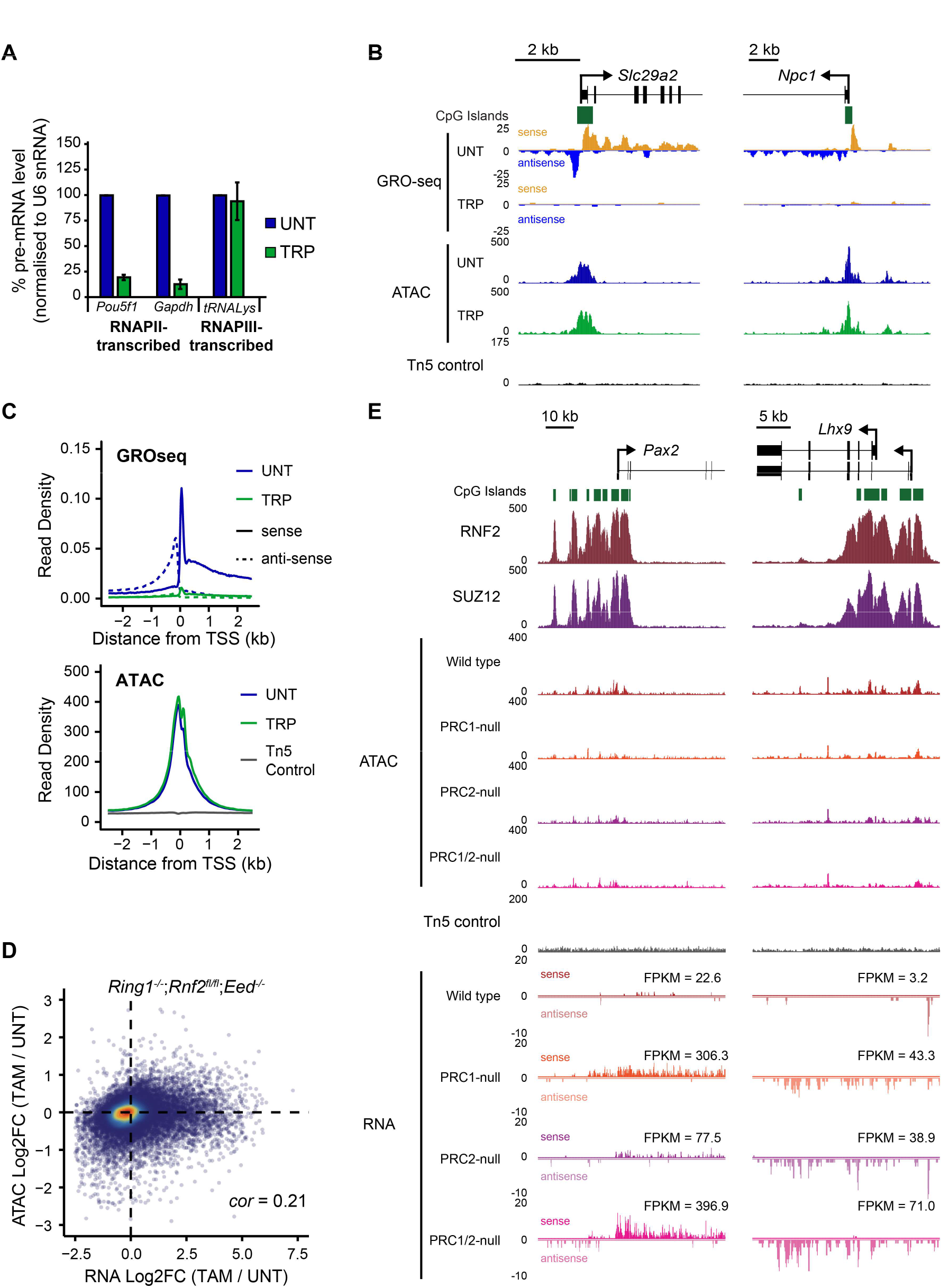
Promoter accessibility is independent of transcriptional activity. A) Quantitative real-time reverse transcriptase PCR for *Pou5f1, Gapdh* and *tRNA-Lys* genes following treatment of mouse E14 ESCs with 500 nM triptolide (TRP) for 50 min, normalised to expression of U6 snRNA and untreated control cells. *n* = 3 ± stdev. B) A genome screenshot at two gene promoters depicting RNA polymerase II engagement (GRO-seq; Jonkers et al. 2014) and ATAC-seq following TRP treatment. C) A metaplot analysis for GRO-seq and ATAC-seq data following TRP treatment at CGI TSS (*n* = 14271). D) A scatterplot comparing the log2 fold change in gene expression (RNA) and chromatin accessibility (ATAC) for CGI promoters in the *Ring1*^-/-^;*Rnf2^fl/fl^*;*Eed*^-/-^ ESCs after tamoxifen (TAM) treatment. E) A genome screenshot for *Ring1*^-/-^;*Rnf2^fl/fl^* and *Ring1*^-/-^;*Rnf2^fl/fl^*;*Eed*^-/-^ ATAC-seq and nuclear RNA-seq signal before and after TAM at two PcG target genes, *Pax2* and *Lhx9*, that are up-regulated after loss of PRC1 and/or PRC2. The normalised expression values (FPKM) for each gene in each cell line and treatment are annotated.

## REFERENCES

Beck S, Lee B-K, Rhee C, Song J, Woo AJ, Kim J. 2014. CpG island-mediated global gene regulatory modes in mouse embryonic stem cells. Nat Commun 5: 5490.

Beh LY, Colwell LJ, Francis NJ. 2012. A core subunit of Polycomb repressive complex 1 is broadly conserved in function but not primary sequence. PNAS 109: E1063–E1071.

Bell O, Schwaiger M, Oakeley EJ, Lienert F, Beisel C, Stadler MB, Schubeler D. 2010. Accessibility of the Drosophila genome discriminates PcG repression, H4K16 acetylation and replication timing. Nat Struct Mol Biol 17: 894–900.

Blackledge NP, Farcas AM, Kondo T, King HW, McGouran JF, Hanssen LL, Ito S, Cooper S, Kondo K, Koseki Y et al. 2014. Variant PRC1 Complex-Dependent H2A Ubiquitylation Drives PRC2 Recruitment and Polycomb Domain Formation. Cell 157: 1445–1459.

Blackledge NP, Klose R. 2011. CpG island chromatin. Epigenetics 6: 147–152.

Blackledge NP, Long HK, Zhou JC, Kriaucionis S, Patient R, Klose RJ. 2012. Bio-CAP: a versatile and highly sensitive technique to purify and characterise regions of non-methylated DNA. Nucleic Acids Res 40: e32–e32.

Blackledge NP, Rose NR, Klose RJ. 2015. Targeting Polycomb systems to regulate gene expression: modifications to a complex story. Nat Rev Mol Cell Biol 16: 643–649.

Boivin A, Dura JM. 1998. In vivo chromatin accessibility correlates with gene silencing in Drosophila. Genetics 150: 1539–1549.

Boyer LA, Plath K, Zeitlinger J, Brambrink T, Medeiros LA, Lee TI, Levine SS, Wernig M, Tajonar A, Ray MK et al. 2006. Polycomb complexes repress developmental regulators in murine embryonic stem cells. Nature 441: 349–353.

Boyle AP, Davis S, Shulha HP, Meltzer P, Margulies EH, Weng Z, Furey TS, Crawford GE. 2008. High-Resolution Mapping and Characterization of Open Chromatin across the Genome. Cell 132: 311–322.

Bracken AP, Dietrich N, Pasini D, Hansen KH, Helin K. 2006. Genome-wide mapping of Polycomb target genes unravels their roles in cell fate transitions. Genes Dev 20: 1123–1136.

Bracken AP, Helin K. 2009. Polycomb group proteins: navigators of lineage pathways led astray in cancer. Nat Rev Cancer 9: 773–784.

Brookes E, de Santiago I, Hebenstreit D, Morris KJ, Carroll T, Xie SQ, Stock JK, Heidemann M, Eick D, Nozaki N et al. 2012. Polycomb associates genome-wide with a specific RNA polymerase II variant, and regulates metabolic genes in ESCs. Cell Stem Cell 10: 157–170.

Buenrostro JD, Giresi PG, Zaba LC, Chang HY, Greenleaf WJ. 2013. Transposition of native chromatin for fast and sensitive epigenomic profiling of open chromatin, DNA-binding proteins and nucleosome position. Nat Methods 10: 1213–1218.

Calabrese JM, Sun W, Song L, Mugford Joshua W, Williams L, Yee D, Starmer J, Mieczkowski P, Crawford Gregory E, Magnuson T. 2012. Site-Specific Silencing of Regulatory Elements as a Mechanism of X Inactivation. Cell 151: 951–963.

Chamberlain SJ, Yee D, Magnuson T. 2008. Polycomb Repressive Complex 2 Is Dispensable for Maintenance of Embryonic Stem Cell Pluripotency. Stem Cells 26: 1496–1505.

Chen K, Xi Y, Pan X, Li Z, Kaestner K, Tyler J, Dent S, He X, Li W. 2013. DANPOS: Dynamic analysis of nucleosome position and occupancy by sequencing. Genome Res 23: 341–351.

Cockerill PN. 2011. Structure and function of active chromatin and DNase I hypersensitive sites. FEBS J 278: 2182–2210.

Cooper S, Dienstbier M, Hassan R, Schermelleh L, Sharif J, Blackledge Neil P, De Marco V, Elderkin S, Koseki H, Klose R et al. 2014. Targeting Polycomb to Pericentric Heterochromatin in Embryonic Stem Cells Reveals a Role for H2AK119u1 in PRC2 Recruitment. Cell Reports 7: 1456–1470.

Cruz-Molina S, Respuela P, Tebartz C, Kolovos P, Nikolic M, Fueyo R, van Ijcken WFJ, Grosveld F, Frommolt P, Bazzi H et al. 2017. PRC2 Facilitates the Regulatory Topology Required for Poised Enhancer Function during Pluripotent Stem Cell Differentiation. Cell Stem Cell 20: 689–705.

de Dieuleveult M, Yen K, Hmitou I, Depaux A, Boussouar F, Dargham DB, Jounier S, Humbertclaude H, Ribierre F, Baulard C et al. 2016. Genome-wide nucleosome specificity and function of chromatin remodellers in ES cells. Nature 530: 113–116.

Deaton AM, Bird A. 2011. CpG islands and the regulation of transcription. Gene Dev 25: 1010–1022.

Deaton AM, Gómez-RodríguezM, MieczkowskiJ, TolstorukovMY, KunduS, SadreyevRI, JansenLET, KingstonRE. 2016. Enhancer regions show high histone H3.3 turnover that changes during differentiation. eLife 5: e15316.

Di Croce L, Helin K. 2013. Transcriptional regulation by Polycomb group proteins. Nat Struct Mol Biol 20: 1147–1155.

Dobin A, Davis CA, Schlesinger F, Drenkow J, Zaleski C, Jha S, Batut P, Chaisson M, Gingeras TR. 2012. STAR: ultrafast universal RNA-seq aligner. Bioinformatics 29: 15–21.

Endoh M, Endo TA, Endoh T, Fujimura Y-i, Ohara O, Toyoda T, Otte AP, Okano M, Brockdorff N, Vidal M et al. 2008. Polycomb group proteins Ring1A/B are functionally linked to the core transcriptional regulatory circuitry to maintain ES cell identity. Development 135: 1513–1524.

Endoh M, Endo TA, Endoh T, Isono K, Sharif J, Ohara O, Toyoda T, Ito T, Eskeland R, Bickmore WA et al. 2012. Histone H2A mono-ubiquitination is a crucial step to mediate PRC1-dependent repression of developmental genes to maintain ES cell identity. PLOS Genet 8: e1002774.

Eskeland R, Leeb M, Grimes GR, Kress C, Boyle S, Sproul D, Gilbert N, Fan Y, Skoultchi AI, Wutz A et al. 2010. Ring1B Compacts Chromatin Structure and Represses Gene Expression Independent of Histone Ubiquitination. Mol Cell 38: 452–464.

Farcas AM, Blackledge NP, Sudbery I, Long HK, McGouran JF, Rose NR, Lee S, Sims D, Cerase A, Sheahan TW et al. 2012. KDM2B links the Polycomb Repressive Complex 1 (PRC1) to recognition of CpG islands. eLife 1: e00205.

Fenouil R, Cauchy P, Koch F, Descostes N, Cabeza JZ, Innocenti C, Ferrier P, Spicuglia S, Gut M, Gut I et al. 2012. CpG islands and GC content dictate nucleosome depletion in a transcription-independent manner at mammalian promoters. Genome Res 22: 2399–2408.

Fitzgerald DP, Bender W. 2001. Polycomb Group Repression Reduces DNA Accessibility. Mol Cell Biol 21: 6585–6597.

Francis NJ, Kingston RE, Woodcock CL. 2004. Chromatin Compaction by a Polycomb Group Protein Complex. Science 306: 1574–1577.

Frank CL, Manandhar D, Gordân R, Crawford GE. 2016. HDAC inhibitors cause site-specific chromatin remodeling at PU.1-bound enhancers in K562 cells. Epigenet Chromatin 9: 1–17.

Gilchrist DA, Dos Santos G, Fargo DC, Xie B, Gao Y, Li L, Adelman K. 2010. Pausing of RNA Polymerase II Disrupts DNA-Specified Nucleosome Organization to Enable Precise Gene Regulation. Cell 143: 540–551.

Giresi PG, Kim J, McDaniell RM, Iyer VR, Lieb JD. 2007. FAIRE (Formaldehyde-Assisted Isolation of Regulatory Elements) isolates active regulatory elements from human chromatin. Genome Res 17: 877–885.

Grau DJ, Chapman BA, Garlick JD, Borowsky M, Francis NJ, Kingston RE. 2011. Compaction of chromatin by diverse Polycomb group proteins requires localized regions of high charge. Gene Dev 25: 2210–2221.

Guertin MJ, Lis JT. 2013. Mechanisms by which transcription factors gain access to target sequence elements in chromatin. Curr Opin Genet Dev 23: 116–123.

Han X, Yu H, Huang D, Xu Y, Saadatpour A, Li X, Wang L, Yu J, Pinello L, Lai S et al. 2017. A molecular roadmap for induced multi-lineage trans-differentiation of fibroblasts by chemical combinations. Cell Research 27: 386.

He J, Shen L, Wan M, Taranova O, Wu H, Zhang Y. 2013. Kdm2b maintains murine embryonic stem cell status by recruiting PRC1 complex to CpG islands of developmental genes. Nat Cell Biol 15: 373–384.

Heinz S, Benner C, Spann N, Bertolino E, Lin YC, Laslo P, Cheng JX, Murre C, Singh H, Glass CK. 2010. Simple Combinations of Lineage-Determining Transcription Factors Prime cis-Regulatory Elements Required for Macrophage and B Cell Identities. Mol Cell 38: 576–589.

Henikoff S. 2008. Nucleosome destabilization in the epigenetic regulation of gene expression. Nat Rev Genet 9: 15–26.

Hinrichs AS, Karolchik D, Baertsch R, Barber GP, Bejerano G, Clawson H, Diekhans M, Furey TS, Harte RA, Hsu F et al. 2006. The UCSC Genome Browser Database: update 2006. Nucleic Acids Res 34: D590–598.

Hodges HC, Stanton BZ, Cermakova K, Chang C-Y, Miller EL, Kirkland JG, Ku WL, Veverka V, Zhao K, Crabtree GR. 2018. Dominant-negative SMARCA4 mutants alter the accessibility landscape of tissue-unrestricted enhancers. Nat Struct Mol Biol 25: 61–72.

Hon GC, Rajagopal N, Shen Y, McCleary DF, Yue F, Dang MD, Ren B. 2013. Epigenetic memory at embryonic enhancers identified in DNA methylation maps from adult mouse tissues. Nat Genet 45: 1198–1206.

Hu G, Schones DE, Cui K, Ybarra R, Northrup D, Tang Q, Gattinoni L, Restifo NP, Huang S, Zhao K. 2011. Regulation of nucleosome landscape and transcription factor targeting at tissue-specific enhancers by BRG1. Genome Res 21: 1650–1658.

Hu X, Zhang L, Mao S-Q, Li Z, Chen J, Zhang R-R, Wu H-P, Gao J, Guo F, Liu W et al. 2014. Tet and TDG Mediate DNA Demethylation Essential for Mesenchymal-to-Epithelial Transition in Somatic Cell Reprogramming. Cell Stem Cell 14: 512–522.

Illingworth RS, Moffat M, Mann AR, Read D, Hunter CJ, Pradeepa MM, Adams IR, Bickmore WA. 2015. The E3 ubiquitin ligase activity of RING1B is not essential for early mouse development. Gene Dev 29: 1897–1902.

Jiang C, Pugh BF. 2009. Nucleosome positioning and gene regulation: advances through genomics. Nat Rev Genet 10: 161–172.

Jonkers I, Kwak H, Lis JT. 2014. Genome-wide dynamics of Pol II elongation and its interplay with promoter proximal pausing, chromatin, and exons. eLife 3: e02407.

Jullien J, Vodnala M, Pasque V, Oikawa M, Miyamoto K, Allen G, David SA, Brochard V, Wang S, Bradshaw C et al. 2017. Gene Resistance to Transcriptional Reprogramming following Nuclear Transfer Is Directly Mediated by Multiple Chromatin-Repressive Pathways. Mol Cell 65: 873–884.e878.

Kadoch C, Williams RT, Calarco JP, Miller EL, Weber CM, Braun SMG, Pulice JL, Chory EJ, Crabtree GR. 2017. Dynamics of BAF–Polycomb complex opposition on heterochromatin in normal and oncogenic states. Nat Genet 49: 213.

Kelly TK, Liu Y, Lay FD, Liang G, Berman BP, Jones PA. 2012. Genome-wide mapping of nucleosome positioning and DNA methylation within individual DNA molecules. Genome Res 22: 2497–2506.

Kent WJ, Sugnet CW, Furey TS, Roskin KM, Pringle TH, Zahler AM, Haussler, David. 2002. The Human Genome Browser at UCSC. Genome Res 12: 996–1006.

King HW, Klose RJ. 2017. The pioneer factor OCT4 requires the chromatin remodeller BRG1 to support gene regulatory element function in mouse embryonic stem cells. eLife 6: e22631.

Kireeva ML, Walter W, Tchernajenko V, Bondarenko V, Kashlev M, Studitsky VM. 2002. Nucleosome Remodeling Induced by RNA Polymerase II: Loss of the H2A/H2B Dimer during Transcription. Mol Cell 9: 541–552.

Kloet SL, Makowski MM, Baymaz HI, van Voorthuijsen L, Karemaker ID, Santanach A, Jansen PWTC, Di Croce L, Vermeulen M. 2016. The dynamic interactome and genomic targets of Polycomb complexes during stem-cell differentiation. Nat Struct Mol Biol 23: 682–690.

Kornberg RD, Lorch Y. 1999. Twenty-Five Years of the Nucleosome, Fundamental Particle of the Eukaryote Chromosome. Cell 98: 285–294.

Kouzarides T. 2007. Chromatin Modifications and Their Function. Cell 128: 693–705.

Ku M, Koche RP, Rheinbay E, Mendenhall EM, Endoh M, Mikkelsen TS, Presser A, Nusbaum C, Xie X, Chi AS et al. 2008. Genomewide Analysis of PRC1 and PRC2 Occupancy Identifies Two Classes of Bivalent Domains. PLOS Genet 4: e1000242.

Kulaeva OI, Hsieh F-K, Studitsky VM. 2010. RNA polymerase complexes cooperate to relieve the nucleosomal barrier and evict histones. PNAS 107: 11325–11330.

Kundu S, Ji F, Sunwoo H, Jain G, Lee JT, Sadreyev RI, Dekker J, Kingston RE. 2017. Polycomb Repressive Complex 1 Generates Discrete Compacted Domains that Change during Differentiation. Mol Cell 65: 432–446.e435.

Langmead B, Salzberg SL. 2012. Fast gapped-read alignment with Bowtie 2. Nat Methods 9: 357–359.

Leeb M, Pasini D, Novatchkova M, Jaritz M, Helin K, Wutz A. 2010. Polycomb complexes act redundantly to repress genomic repeats and genes. Gene Dev 24: 265–276.

Lehmann L, Ferrari R, Vashisht AA, Wohlschlegel JA, Kurdistani SK, Carey M. 2012. Polycomb Repressive Complex 1 (PRC1) Disassembles RNA Polymerase II Preinitiation Complexes. J Biol Chem 287: 35784–35794.

Lennartsson A, Arner E, Fagiolini M, Saxena A, Andersson R, Takahashi H, Noro Y, Sng J, Sandelin A, Hensch TK et al. 2015. Remodeling of retrotransposon elements during epigenetic induction of adult visual cortical plasticity by HDAC inhibitors. Epigenet Chromatin 8: 1–15.

Li B, Carey M, Workman JL. 2007. The Role of Chromatin during Transcription. Cell 128: 707–719.

Li H, Handsaker B, Wysoker A, Fennell T, Ruan J, Homer N, Marth G, Abecasis G, Durbin R. 2009. The Sequence Alignment/Map format and SAMtools. Bioinformatics 25: 2078–2079.

Liang Z, Brown KE, Carroll T, Taylor B, Vidal IF, Hendrich B, Rueda D, Fisher AG, Merkenschlager M. 2017. A high-resolution map of transcriptional repression. eLife 6: e22767.

Long HK, Sims D, Heger A, Blackledge NP, Kutter C, Wright ML, Grützner F, Odom DT, Patient R, Ponting CP et al. 2013. Epigenetic conservation at gene regulatory elements revealed by nonmethylated DNA profiling in seven vertebrates. eLife 2: e00348.

Love MI, Huber W, Anders S. 2014. Moderated estimation of fold change and dispersion for RNA-seq data with DESeq2. Genome Biol 15: 1–21.

Marathe HG, Mehta G, Zhang X, Datar I, Mehrotra A, Yeung KC, de la Serna IL. 2013. SWI/SNF Enzymes Promote SOX10- Mediated Activation of Myelin Gene Expression. PLOS ONE 8: e69037.

Margueron R, Li G, Sarma K, Blais A, Zavadil J, Woodcock CL, Dynlacht BD, Reinberg D. 2008. Ezh1 and Ezh2 Maintain Repressive Chromatin through Different Mechanisms. Mol Cell 32: 503–518.

McCall K, Bender W. 1996. Probes of chromatin accessibility in the Drosophila bithorax complex respond differently to Polycomb-mediated repression. EMBO J 15: 569–580.

Mendenhall EM, Koche RP, Truong T, Zhou VW, Issac B, Chi AS, Ku M, Bernstein BE. 2010. GC-Rich Sequence Elements Recruit PRC2 in Mammalian ES Cells. PLOS Genet 6: e1001244.

Mieczkowski J, Cook A, Bowman SK, Mueller B, Alver BH, Kundu S, Deaton AM, Urban JA, Larschan E, Park PJ et al. 2016. MNase titration reveals differences between nucleosome occupancy and chromatin accessibility. Nat Commun 7: 11485.

Mueller B, Mieczkowski J, Kundu S, Wang P, Sadreyev R, Tolstorukov MY, Kingston RE. 2017. Widespread changes in nucleosome accessibility without changes in nucleosome occupancy during a rapid transcriptional induction. Gene Dev 31: 451–462.

Muller J, Verrijzer P. 2009. Biochemical mechanisms of gene regulation by polycomb group protein complexes. Curr Opin Genet Dev 19: 150–158.

Norton VG, Imai BS, Yau P, Bradbury EM. 1989. Histone acetylation reduces nucleosome core particle linking number change. Cell 57: 449–457.

Pengelly AR, Kalb R, Finkl K, Muller J. 2015. Transcriptional repression by PRC1 in the absence of H2A monoubiquitylation. Genes Dev 29: 1487–1492.

Picelli S, Björklund ÅK, Reinius B, Sagasser S, Winberg G, Sandberg R. 2014. Tn5 transposase and tagmentation procedures for massively scaled sequencing projects. Genome Res 24: 2033–2040.

Quinlan AR. 2014. BEDTools: The Swiss-Army Tool for Genome Feature Analysis. Curr Protoc Bioinformatics 47: 11.12.11–11.12.34.

Ramírez J, Dege C, Kutateladze TG, Hagman J. 2012. MBD2 and Multiple Domains of CHD4 Are Required for Transcriptional Repression by Mi-2/NuRD Complexes. Mol Cell Biol 32: 5078–5088.

Ran FA, Hsu PD, Wright J, Agarwala V, Scott DA, Zhang F. 2013. Genome engineering using the CRISPR-Cas9 system. Nat Protoc 8: 2281–2308.

Reynolds N, Salmon-DivonM, DvingeH, Hynes-AllenA, BalasooriyaG, LeafordD, BehrensA, BertoneP, HendrichB. 2012. NuRD‐mediated deacetylation of H3K27 facilitates recruitment of Polycomb Repressive Complex 2 to direct gene repression. EMBO J 31: 593–605.

Rincon-Arano H, Halow J, Delrow Jeffrey J, Parkhurst Susan M, Groudine M. 2012. UpSET Recruits HDAC Complexes and Restricts Chromatin Accessibility and Acetylation at Promoter Regions. Cell 151: 1214–1228.

Rose NR, King HW, Blackledge NP, Fursova NA, Ember KJI, Fischer R, Kessler BM, Klose RJ. 2016. RYBP stimulates PRC1 to shape chromatin-based communication between Polycomb repressive complexes. eLife 5: e18591.

Schep AN, Buenrostro JD, Denny SK, Schwartz K, Sherlock G, Greenleaf WJ. 2015. Structured nucleosome fingerprints enable high-resolution mapping of chromatin architecture within regulatory regions. Genome Res 25: 1757–1770.

Schoenfelder S, Sugar R, Dimond A, Javierre B-M, Armstrong H, Mifsud B, Dimitrova E, Matheson L, Tavares-Cadete F, Furlan-Magaril M et al. 2015. Polycomb repressive complex PRC1 spatially constrains the mouse embryonic stem cell genome. Nat Genet 47: 1179–1186.

Shen X, Liu Y, Hsu YJ, Fujiwara Y, Kim J, Mao X, Yuan GC, Orkin SH. 2008. EZH1 Mediates Methylation on Histone H3 Lysine 27 and Complements EZH2 in Maintaining Stem Cell Identity and Executing Pluripotency. Mol Cell 32: 491–502.

Shogren-Knaak M, Ishii H, Sun J-M, Pazin MJ, Davie JR, Peterson CL. 2006. Histone H4-K16 Acetylation Controls Chromatin Structure and Protein Interactions. Science 311: 844–847.

Simon Jeffrey A, Kingston Robert E. 2013. Occupying Chromatin: Polycomb Mechanisms for Getting to Genomic Targets, Stopping Transcriptional Traffic, and Staying Put. Mol Cell 49: 808–824.

Song L, Zhang Z, Grasfeder LL, Boyle AP, Giresi PG, Lee B-K, Sheffield NC, Gräf S, Huss M, Keefe D et al. 2011. Open chromatin defined by DNaseI and FAIRE identifies regulatory elements that shape cell-type identity. Genome Res 21: 1757–1767.

Stark R, Brown G. 2011. DiffBind: differential binding analysis of ChIP-Seq peak data. In Bioconductor, http://bioconductor.org/packages/release/bioc/vignettes/DiffBind/inst/doc/DiffBind.pdf

Stock JK, Giadrossi S, Casanova M, Brookes E, Vidal M, Koseki H, Brockdorff N, Fisher AG, Pombo A. 2007. Ring1-mediated ubiquitination of H2A restrains poised RNA polymerase II at bivalent genes in mouse ES cells. Nat Cell Biol 9: 1428–1435.

Tavares L, Dimitrova E, Oxley D, Webster J, Poot R, Demmers J, Bezstarosti K, Taylor S, Ura H, Koide H et al. 2012. RYBP-PRC1 Complexes Mediate H2A Ubiquitylation at Polycomb Target Sites Independently of PRC2 and H3K27me3. Cell 148: 664–678.

Thakurela S, Garding A, Jung J, Schübeler D, Burger L, Tiwari VK. 2013. Gene regulation and priming by topoisomerase IIα in embryonic stem cells. Nat Commun 4: 2478.

Thurman RE, Rynes E, Humbert R, Vierstra J, Maurano MT, Haugen E, Sheffield NC, Stergachis AB, Wang H, Vernot B et al. 2012. The accessible chromatin landscape of the human genome. Nature 489: 75–82.

Trojer P, Cao Alina R, Gao Z, Li Y, Zhang J, Xu X, Li G, Losson R, Erdjument-Bromage H, Tempst P et al. 2011. L3MBTL2 Protein Acts in Concert with PcG Protein-Mediated Monoubiquitination of H2A to Establish a Repressive Chromatin Structure. Mol Cell 42: 438–450.

Trojer P, Li G, Sims Iii RJ, Vaquero A, Kalakonda N, Boccuni P, Lee D, Erdjument-Bromage H, Tempst P, Nimer SD et al. 2007. L3MBTL1, a Histone-Methylation-Dependent Chromatin Lock. Cell 129: 915–928.

Ura H, Usuda M, Kinoshita K, Sun C, Mori K, Akagi T, Matsuda T, Koide H, Yokota T. 2008. STAT3 and Oct-3/4 Control Histone Modification through Induction of Eed in Embryonic Stem Cells. J Biol Chem 283: 9713–9723.

West JA, Cook A, Alver BH, Stadtfeld M, Deaton AM, Hochedlinger K, Park PJ, Tolstorukov MY, Kingston RE. 2014. Nucleosomal occupancy changes locally over key regulatory regions during cell differentiation and reprogramming. Nat Commun 5: 4719.

Whyte WA, Bilodeau S, Orlando DA, Hoke HA, Frampton GM, Foster CT, Cowley SM, Young RA. 2012. Enhancer decommissioning by LSD1 during embryonic stem cell differentiation. Nature 482: 221.

Wu X, Johansen JV, Helin K. 2013. Fbxl10/Kdm2b recruits polycomb repressive complex 1 to CpG islands and regulates H2A ubiquitylation. Mol Cell 49: 1134–1146.

Yildirim O, Li R, Hung J-H, Chen P, Dong X, Ee L-S, Weng Z, Rando O, Fazzio T. 2011. Mbd3/NURD Complex Regulates Expression of 5-Hydroxymethylcytosine Marked Genes in Embryonic Stem Cells. Cell 147: 1498–1510.

YueF ChengY BreschiA VierstraJ WuW RybaT SandstromR MaZ DavisC PopeBD et al. 2014. A comparative encyclopedia of DNA elements in the mouse genome. Nature 515: 355–364.

Zhang Y, Liu T, Meyer CA, Eeckhoute J, Johnson DS, Bernstein BE, Nusbaum C, Myers RM, Brown M, Li W et al. 2008. Model-based Analysis of ChIP-Seq (MACS). Genome Biol 9: 1–9.

Zink D, Paro R. 1995. Drosophila Polycomb-group regulated chromatin inhibits the accessibility of a trans-activator to its target DNA. EMBO J 14: 5660–5671.

